# Systemic resistance to pathogens in Arabidopsis requires HASTY-dependent miRNA cell-to-cell movement

**DOI:** 10.1101/2025.10.02.680033

**Authors:** Musso Manuel, Nahir Alanie, Quevedo Luciano, Trenchi Alejandra, Cecchini Nicolás Miguel, Lascano Hernán Ramiro, Cambiagno Damian Alejandro

## Abstract

Plant defenses against pathogens are tightly regulated through complex gene expression control mechanisms. The precise activation and repression of defense-related genes are crucial to balancing the trade-off between growth and immunity. Micro RNAs (miRNAs) play a well-established role in the local regulation of plant-microbe interactions. While some miRNAs are also essential for systemic defense responses, their mechanisms of action, biogenesis, and long-distance mobility remain largely unexplored. Here, we show that HASTY (HST), a key factor in miRNA biogenesis and intercellular movement, is required for systemic defense activation. The impaired mobility of miRNAs in *hst* mutants correlates with a lack of systemic responses. In infected tissues, HST may enhance the co-transcriptional processing of specific pri-miRNAs, which promotes the cell-to-cell movement of their mature miRNAs and contributes to the activation of systemic defenses. Furthermore, two miRNAs that exhibit increased mobility during systemic defense induction are required for a proper systemic response. Interestingly, complementing *hst* mutants with a version of HST expressed exclusively in companion cells is sufficient to restore systemic defense induction, highlighting the role of miRNA cell-to-cell movement. These findings shed light on the role of HST in plant immunity, linking miRNA biogenesis and mobility to the fine-tuned regulation of systemic defenses.

## Introduction

Recognition of pathogen-associated molecular patterns (PAMPs) or pathogen effectors by extracellular and/or intracellular receptors, induce a defense signaling cascade involving MAPKs, reactive oxygen species, and the accumulation of hormones such as salicylic acid (SA) (Jones et al., 2016; Macho and Zipfel, 2014; Weralupitiya et al., 2024). Notably, resistance induction occurs either in cells perceiving pathogens or systemically in neighboring and distal tissues. Among the best-characterized systemic resistance program is the systemic acquired resistance (SAR) (Conrath et al., 2015). This systemic defense is mediated by mobile signals originating from the infected site, including non-protein amino acids like pipecolic acid (PIP) and N-hydroxy pipecolic acid (NHP), hormones, and volatile compounds (Vlot et al., 2020). The exogenous treatments with PIP/NHP have been shown to trigger both local and systemic defenses, with these signals being transported through the phloem, and accessing it via phloem companion cells (CCs) (Foret et al., 2024; Vlot *et al*., 2020; Yildiz et al., 2021).

Several miRNAs have been identified as key regulators of plant immune responses (Asadi and Millar, 2024). Their importance is highlighted by studies showing that mutations in components of the miRNA biogenesis pathway impair plant immunity (Pelaez and Sanchez, 2013). Moreover, multiple components of this pathway are targeted by pathogen effectors (Parperides et al., 2023). miRNA biogenesis begins with the transcription of *MIR* genes by RNA Polymerase II, generating single-stranded RNAs that fold into stem-loop structures. These primary miRNAs (pri-miRNAs) are processed by a miRNA biogenesis complex, which includes the endonuclease Dicer-Like 1 (DCL1) and cofactors such as SERRATE (SE) and HYLAROUS 1 (HYL1) (Achkar et al., 2016). Notably, DCL1 can function at the co-transcriptional level, being recruited to chromatin by the karyopherin HASTY (HST), the Mediator Complex subunit 37, and Elongator complex (Cambiagno et al., 2021; Fang et al., 2015). This co-transcriptional processing relies on R-loops that facilitate the interaction between nascent pri-miRNAs and *MIR* loci. In contrast, miRNAs whose biogenesis is independent of HST appear to be processed post-transcriptionally (Gonzalo et al., 2022).

While most research has focused on how miRNAs regulate local plant immune responses (Song et al., 2021), their role in systemic defenses remains less understood. miR390 is required for SAR induction, facilitating the production of AGO7-dependent trans-acting small interfering RNAs (tasiRNAs) D7 and D8 from the TAS3a transcript. These tasiRNAs move to distal tissues, where they are loaded into AGO1 and target auxin response factor mRNAs (Shine et al., 2022). miRNA2111 was identified as a systemic regulator of plant-microbe interactions in the symbiotic relationship between legumes and rhizobia (Tsikou et al., 2018). miR2111 moves from the shoot to the root, promoting nodulation by repressing the expression of the symbiosis suppressor TOO MUCH LOVE (Tsikou *et al*., 2018). Both tasiRNAs D7/D8 and miR2111 act in a non-cell autonomous manner. In this case, miRNA biogenesis occurs in a source cell, and the miRNAs spread to neighboring or distal cells either through cell-to-cell movement or via the phloem, ultimately regulating their targets in a sink tissue (Shine *et al*., 2022; Tsikou *et al*., 2018). Additionally, plant miRNAs can modulate infection by directly silencing pathogen virulence effectors. Those miRNAs are known as trans-kingdom miRNAs, moving from plants to fungal or oomycete pathogens via extracellular vesicles (EVs) (Cai et al., 2023).

The systemic movement of miRNAs differs from that of other RNAs (Li et al., 2021; Park et al., 2005). Systemic miRNA movement via cell-to-cell transport appears to be inhibited by their association with AGO1, as AGO-free small RNAs (sRNAs) are more likely to move between cells (Devers et al., 2020). Recently, it was shown that HST positively regulates the movement of non-cell-autonomous miRNAs (Brioudes et al., 2021). HST interacts with the MEDIATOR (MED) complex and DCL1, promoting *MIR* gene co-transcriptional processing at the nuclear pore complex (NPC) (Cambiagno *et al*., 2021; Gonzalo et al., 2025). This localization near the NPC may facilitate the direct translocation of miRNAs to the cytoplasm, bypassing AGO1 loading and promoting their movement between cells. In contrast, HAWAIIAN SKIRT (HWS) targets MED subunits at the NPC for degradation, disrupting the MIR-HST-NPC interaction and miRNA distal movement, affecting co-transcriptional pri-miRNA processing in this nuclear region (Gonzalo *et al*., 2025). Additionally, the microtubule-severing enzyme subunit KATANIN1 inhibits miRNA loading into AGO1 in source cells, preventing their non-cell-autonomous activity in distal cells (Fan et al., 2022).

Although miRNA role in plant defenses was extensively studied, little is known about the mechanisms regulating miRNA biogenesis, mobility, and activity during systemic defense induction. Here, we show that HST is essential for systemic defense induction, probably by promoting cell-to-cell movement of miRNAs. In support of this, systemic defense responses can be restored in the *hst* mutant either by rescuing miRNA mobility or by complementing HST expression specifically in CCs. We observed that systemic defense activation promotes a shift toward co-transcriptional processing of miRNA precursors, particularly for those miRNAs with enhanced cell-to-cell mobility. Furthermore, HST subcellular localization is altered upon treatment with the mobile signal PIP. Accordingly, mutants that affect HST subcellular localization are defective in systemic defenses induction. These findings position HST as a central regulator of local and systemic immune responses, underscoring the intricate control of miRNA biogenesis and mobility during the activation of pathogen-induced defenses.

## Results

### HASTY differentially modulates local and systemic defenses

Among small RNAs, miRNAs play a key role in regulating plant defense responses, as evidenced by altered systemic resistance and defense activation in mutants defective in miRNA biogenesis (Rebolledo-Prudencio et al., 2021; Silvestri et al., 2024). Given HASTY’s (HST) role in miRNA processing and movement (Brioudes *et al*., 2021; Cambiagno *et al*., 2021), we evaluated its function in local and systemic defenses. For this, we analyzed susceptibility to *Pseudomonas cannabina* pv. *alisalensis* [formerly called *P. syringae* pv. *maculicola* ES4326 (*Pma*) (Baltrus et al., 2011; Sarris et al., 2013)] in plants pre-infected in the lower leaves two days earlier with or without *P. syringae pv. tomato* (*Pst*)-*AvrRpt2* to induce systemic resistance (SAR) or assess local susceptibility. We found that *hst-15* plants exhibited reduced susceptibility to *Pma* compared to WT plants under mock treatment of the lower leaves (Fig. 1A), suggesting that *hst-15* has increased local resistance to pathogens. In contrast, when comparing *Pma* growth between mock-treated and SAR-induced plants, we found that *hst-15* mutants failed to fully induce SAR (Fig. 1A-B). We further tested the ability of *hst-15* and *hst-3* allelic mutants to induce systemic resistance upon watering plants with the mobile signal PIP. Both mutants exhibited increased resistance under mock-treated conditions, although to different extents (Fig. 1C), yet neither *hst-15* nor *hst-3* induced a significant systemic resistance to *Pma* compared to WT plants (Fig. 1C-D).

**Figure 1.**
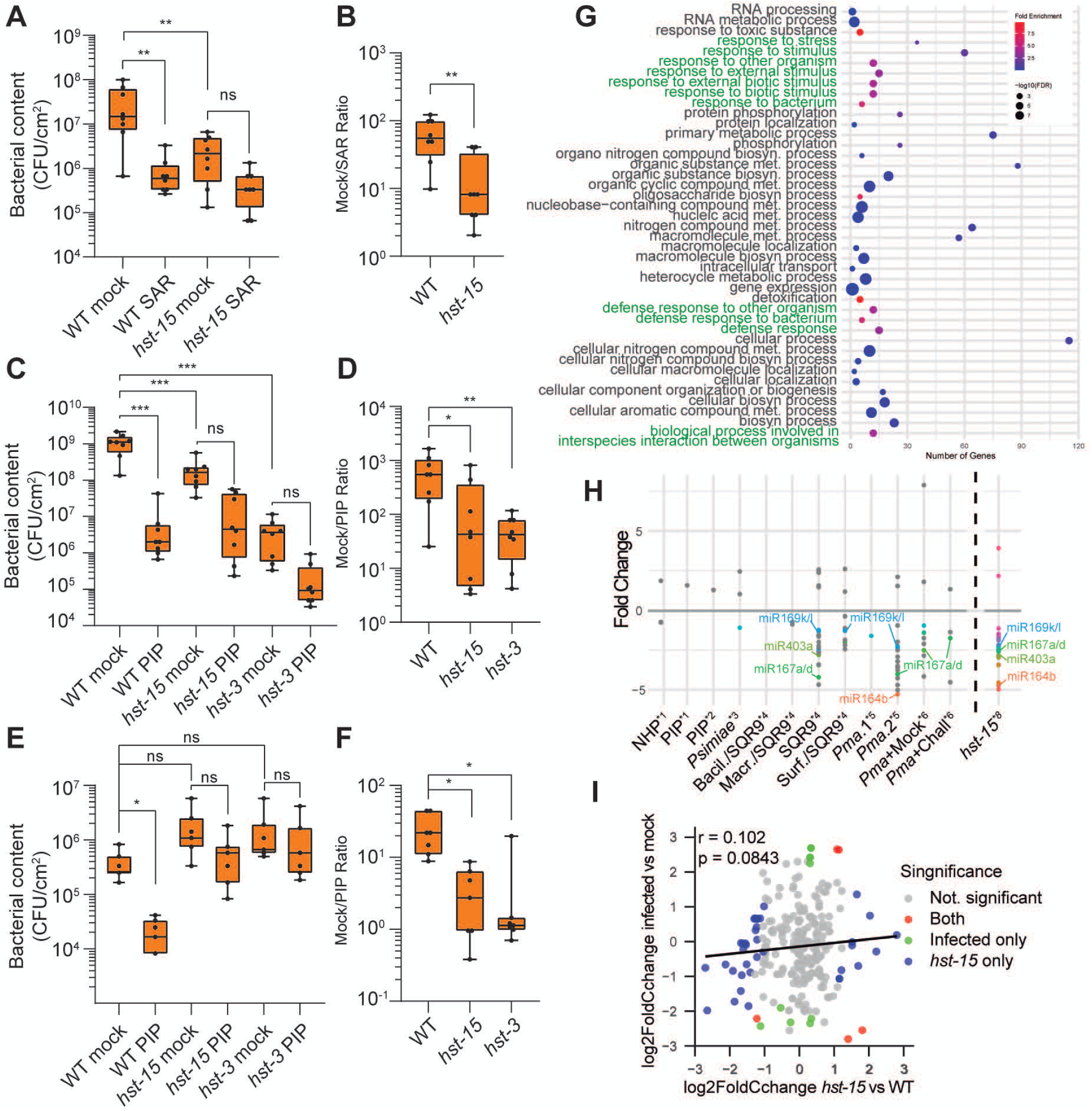
Local and systemic defense responses in hst mutants. (A, B) Growth of *Pma* in distal leaves of wild-type (WT) and *hst-15* plants pre-treated systemically with mock or *Pst-AvrRpt2* for SAR. (C-F) Growth of *Pma* in distal leaves after systemic mock or PIP treatment. Inoculation with *Pma* was performed by syringe infiltration (C, D) or spray (E, F). Pathogen growth is shown as CFU/cm^2^ (A, C, E) or as the ratio between mock and SAR/PIP treatments (B, D, **F).** Statistical significance (*p < 0.05; **p < 0.01; ***p < 0.001) was evaluated using ANOVA, with multiple comparisons corrected using Dunn’s test (A, C-F), or two-tailed unpaired t-test (B). (G) Dot plot of GO-term analysis showing enriched biological processes among upregulated genes in non-infected *hst-15* plants. Pathogen-related terms are highlighted in green. **(H)** Dot plot of differentially expressed MIR genes in tissues with systemically induced defenses. Data were obtained from six transcriptomes [*1: (Yildiz et al., 2021); *2: (Hartmann et al., 2018); *3: (Desrut et al., 2020); *4: (SRP258528); *5: (Bernsdorff et al., 2016); *6: (Baum et al., 2019)]. The dark dashed line separates these datasets from misregulated miRNAs in non-treated *hst-15* plants [*8: (Cambiagno et al., 2021)]. All miRNAs misregulated in *hst-15* are highlighted in color, and the six whose *MIR* expression is also affected during systemic defense are labeled. (I) Scatter plot comparing miRNA expression changes in *hst-15* and *Pst-AvrRpt2-infected* plants. Dots indicate miRNAs that are not significantly different from their controls (grey), misregulated in both samples (red), or significant only in infected plants (green) or in *hst-15* (blue). Pearson correlation (r) and p-value are shown in the upper-left corner.

Similarly, systemic defense induction was not observed in *hst* mutants following spray inoculation with *Pma* (Fig. 1E–F), although susceptibility to the pathogen under mock treatment was comparable to WT. This suggests that HST contributes to systemic defense and may modulate plant responses to infiltration-related stress. Consistently, GO-term analysis of upregulated genes in *hst-15* showed enrichment in biotic stress– related categories (Fig. 1G), possibly reflecting mild stress during growth conditions previously reported (Cambiagno *et al*., 2021). Moreover, there is partial overlap between miRNAs differentially accumulated in *hst-15*, their target mRNA abundance, and *MIR* genes differentially regulated during systemic defense activation (Fig. 1H, S1, see Material and Methods). Unexpectedly, the miRNA profile in infected plants showed no correlation with that of *hst-15* (Fig. 1I). Thus, while changed miRNAs may account for the increased resistance of *hst-15*, they do not explain the impaired systemic response.

### Increased cell-to-cell movement of amiRNAs during systemic defense activation

One possibility is that impaired systemic defense activation in *hst-15* may be linked to the distal movement of miRNAs during stress. If so, miRNA cell-to-cell movement might be affected during the induction of this program in wild-type plants. To test this, we assessed miRNA movement in locally infected leaves to induce SAR. We used *pSUC2:amiR-SUL* transgenic plants, a system previously used to study HST-dependent miRNA cell-to-cell mobility (Brioudes *et al*., 2021; Gonzalo *et al*., 2025). These plants express an artificial miRNA (amiRSUL) in companion cells that silences the *SULFUR* gene, causing chlorosis in both companion and neighboring mesophyll cells. Variations in miRNA movement are reflected in the size of the chlorotic area (de Felippes et al., 2011). Leaves from *pSUC2:amiR-SUL* transgenic plants were locally treated with *Pst-AvrRpt2* or *Pma* to induce SAR, and chlorotic areas were compared to mock-treated controls 24 hours post-treatment. Our results indicate that both *Pst-AvrRpt2* and *Pma* enhance amiRSUL mobility (Fig. 2A). In line with this, infected leaves showed higher yellow intensity in chlorotic areas than non-infected leaves (Fig. 2B). These results indicate that miRNA movement may be modulated by local infection, which could promote their subsequent phloem loading and systemic spread during SAR.

**Figure 2:**
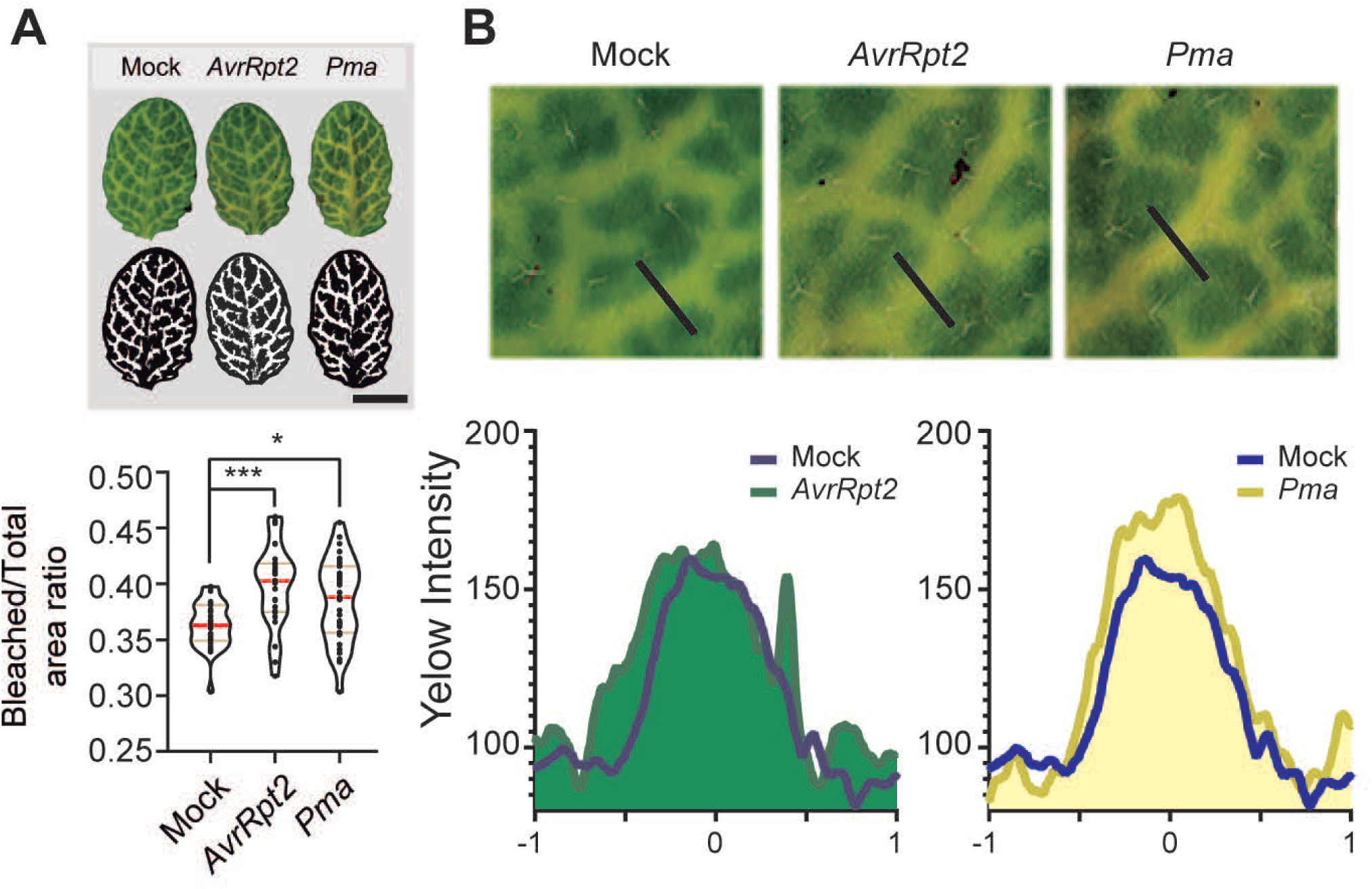
Cell-to-cell movement of artificial miRNAs during defense induction. (A) Leaves from *pSUC2:amiRSUL* transgenic plants untreated (Mock), or treated with *Pst-AvrRpt2 (AvrRpt2)* or *Pma.* Upper panel shows representative images of leaves and corresponding binary masks used to quantify bleached areas. Lower panel: violin plot showing the ratio of bleached area (around vascular tissue) to total leaf area for each treatment. Scale bar represent 0.5 cm. (B) Representative images of leaf veins from untreated (Mock) or infected *(Pst-AvrRpt2* or Pma) plants. A line was drawn along the indicated vein region (black line), and the corresponding yellow intensity profile was extracted.

### Systemic defense induction triggers long-distance movement of endogenous miRNAs

Considering the above result, we next aimed to identify endogenous miRNAs that could be transported from infected tissues during systemic defense activation. To this end, we performed an *in-silico* analysis of RNA-seq datasets from different experiments analyzing local and systemic tissues of infected plants (Baum et al., 2019; Bernsdorff et al., 2016; Desrut et al., 2020; Hartmann et al., 2018; Stringlis et al., 2018; Yildiz *et al*., 2021). We compared pri-miRNA levels to identify potential miRNAs that act as markers of systemic defenses. Among the twenty-one samples analyzed, four pri-miRNAs stood out by showing altered abundance in nine to ten samples, while the rest were affected in five or fewer (Fig. S2A). Pri-miR163 was upregulated, whereas pri-miR414, pri-miR159b, and pri-miR822a were downregulated (Fig. S2A). Notably, except for miR414, all these miRNAs have been previously identified in petiole exudates of untreated plants (Bakirbas et al., 2023), supporting our approach and strongly suggesting their potential role as systemic defense markers.

Thus, we evaluated the abundance of these miRNAs during systemic defense induction in WT plants by pre-treating with PIP followed by distal leaf infection with or without *Pma*. RT-qPCR analysis revealed accumulation of miR163, no change in miR414 or miR822, and a slight reduction in miR159 in *Pma*-infected leaves, both with and without prior PIP treatment (Fig. S2B). We then assessed whether the levels of these miRNAs in phloem exudates from *Pma*-infected leaves (Pathogen Exudates, PEX) differed from those in mock-treated plants (Mock Exudates, MEX) (Fig. 3A). Interestingly, we observed an increased abundance in PEX for three endogenous miRNAs identified as potential systemic defense markers: miR163, miR414, and miR822 (Fig. 3B). In case of miR163, this accumulation could be related to its increased abundance in total tissue (Fig. S2B). We also tested whether these endogenous miRNAs accumulated in phloem exudates from PIP-treated tissues (PIPEX) (Fig. 3A), a strong inducer of local and systemic defenses. All four miRNAs (miR414, miR159b, miR163, and miR822) increased in PIPEX, despite no changes in total leaf tissue (Fig. 3C). As a positive control, we included miR390 and its mobile product tasi-RNA-D7, which accumulated in PIPEX as expected (Shine *et al*., 2022). In contrast, miR171c, miR402, and ACTIN mRNA—a marker of cellular damage—were undetectable in these samples (Fig. S2C) (Bakirbas *et al*., 2023).

**Figure 3:**
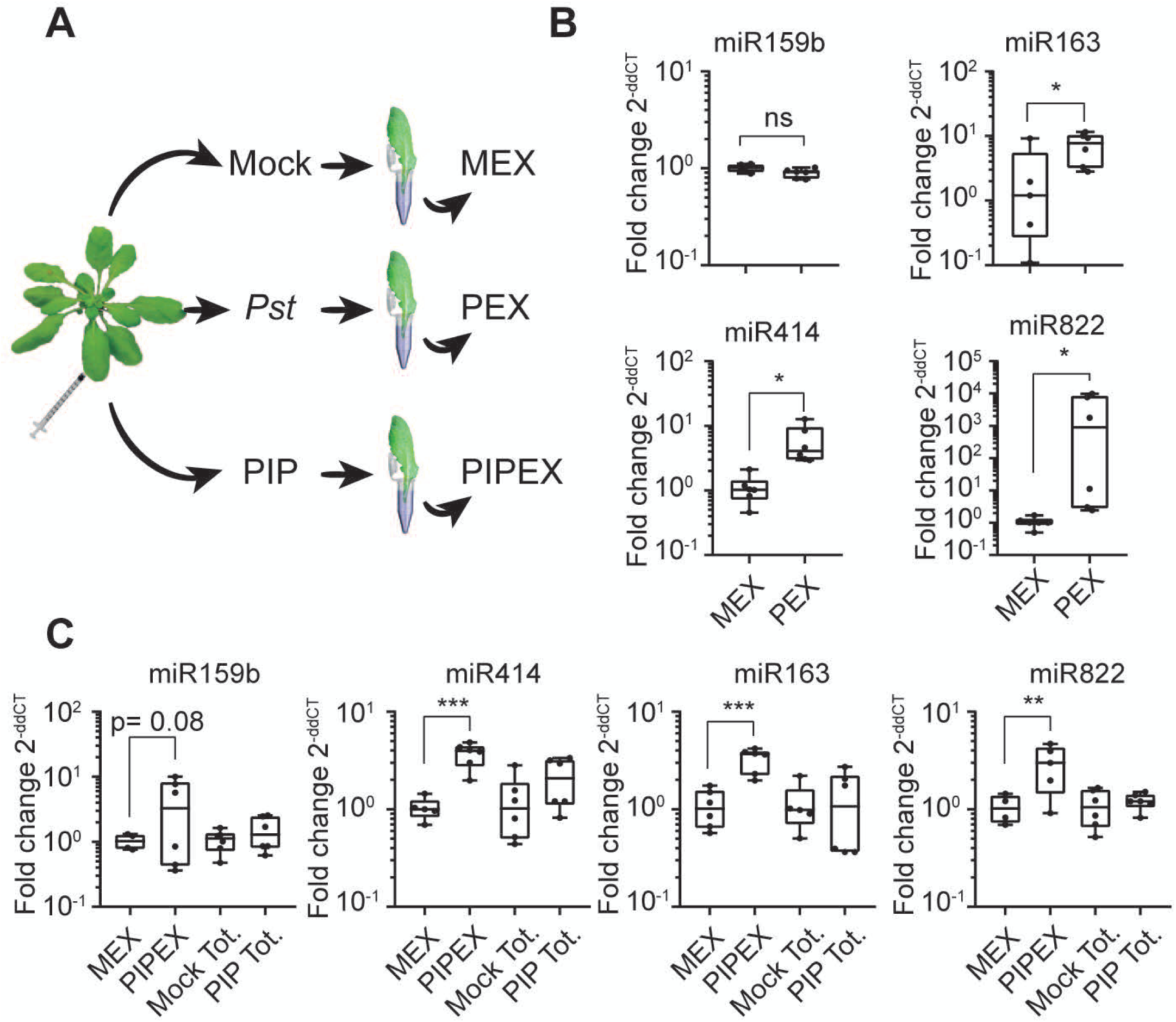
Systemic movement of endogenous miRNAs during defense induction: (A) Scheme of samples coming from phloem exudates from Mock (MEX), Pma-infected (PEX) or PIP-treated (PIPEX) leaves. (B, C) miRNA abundance measured by RT-qPCR in MEX, PEX (B), or PIPEX (C) in wild-type leaves. *: p < 0.05, **: p < 0.01, ***: p < 0.001 by two-tailed unpaired !-test or Mann-Whitney test (B) or by ANOVA with multiple comparisons corrected using Dunn’s test (C).

Together, these results suggest that the long-distance movement of certain miRNAs is promoted in tissues, especially phloem-associated ones, that trigger systemic defense signaling.

### Systemic defense activation favors co-transcriptional processing of mobile miRNAs

HST mutation impairs full induction of systemic defenses (Fig. 1). The non-cell-autonomous activity of many miRNAs relies on co-transcriptional processing, which involves HST and the NPC (Cambiagno *et al*., 2021; Gonzalo *et al*., 2025; Gonzalo *et al*., 2022). Thus, the increased distal movement of specific miRNAs during systemic defense induction may be associated with their mode of processing. To test this, we isolated chromatin-bound and nucleoplasmic pri-miRNAs from leaves treated or not with the systemic resistance inducer PIP. In those samples, we measured the ratio of processed to unprocessed precursors for miR159, miR163, and miR822, using miR171 as a negative control. We found that the ratio of processed to unprocessed precursors for miR159, miR163, and miR822 was higher at chromatin in PIP-treated leaves compared to mock. In contrast, no differences were observed in the nucleoplasm (Fig. 4A). The miR171 showed no difference either at chromatin or nucleoplasm, as expected.

**Figure 4:**
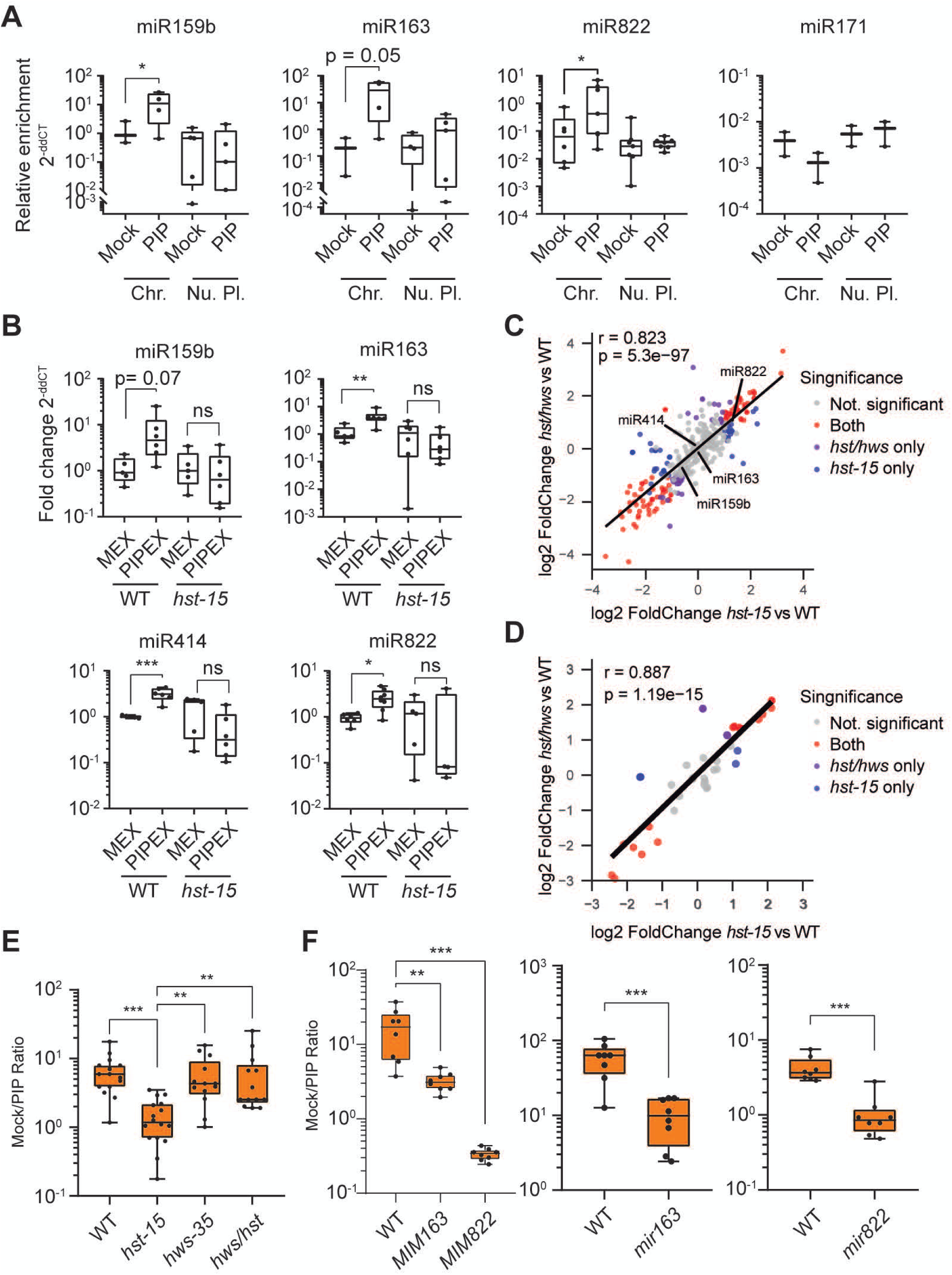
Co/post-transcriptional processing of pri-miRNAs, HST-dependent systemic movement, and requirement of specific miRNAs for systemic defenses. (A) Co-and post-transcriptional processing of selected pri-miRNAs in chromatin-or nucleoplasm-enriched fractions from Mock-or PIP-treated leaves, measured by RT-qPCR. (B) miRNA abundance measured by RT-qPCR in phloem exudates collected from Mock-treated (MEX) or PIP-treated (PIPEX) leaves of wild-type (WT) and *hst-15* plants. (C, D) Scatter plot comparing the abundance of global (C) or defense responsive (D) miRNA in *hst-15* and *hws/hst.* Dots indicate miRNAs that are not significantly different from their controls (grey), misregulated in both samples (red), or significant only in *hws/hst* {purple) or in *hst-15* (blue). Pearson correlation (r) and p-value are shown in the upper-left corner. (E, F) Ratio of pathogen growth between pre-treated systemically with mock or PIP in *hst-15, hws-35*, and *hws-35/hst-15* double mutant *(hws/hst)* (C), or *MIM163, MIM822, mir163*, and *mir822* lines (D). *: p < 0.05, **: p < 0.01, ***: p < 0.001 by ANOVA (A, B, D) or Kruskal-Wallis test (C) with multiple comparisons corrected using Dunn’s test; or by two-tailed unpaired t-test or Mann-Whitney test (D).

These results indicate that the three miRNAs are preferentially processed co-transcriptionally during systemic defense induction, a process correlated with their increased distal movement and suggesting a role for HST.

### HST mediates miRNA distal movement in systemic defense induction

We next tested whether HST affects the increased distal movement of miRNAs during systemic defense induction. We measured endogenous marker miRNAs in MEX and PIPEX from wild-type and *hst-15* mutant. Notably, unlike WT, the *hst* mutation abolished miRNAs accumulation in PIPEX (Fig. 4B), strongly suggesting that HST plays a central role in this process. Negative controls miR171 and Actin were undetected as expected (Fig. S3).

Because HST promotes pri-miRNA processing and movement while HWS represses it, and HWS mutation restores mobility in *hst-15* (*hws-35/hst-15* double mutant) without changing overall miRNA abundance (Gonzalo *et al*., 2025), we tested miRNA levels and confirmed that miR159b, miR163, miR414, and miR822 do not differ between *hst-15* and *hws/hst* double mutant[(Gonzalo *et al*., 2025), Fig. 4C]. Likewise, miRNAs differentially expressed in infected plants [(Zhang et al., 2011a), see Materials and Methods] show no abundance changes between these mutants (Fig. 4D). Moreover, miR159b, miR163, miR414, and miR822 accumulated in the phloem of WT but not *hst-15*, and this accumulation was partially restored in the *hws/hst* double mutant (Fig. S5). Given these results, we tested whether restoring miRNA mobility in the *hst-15* background also restores systemic defenses. WT, *hst-15*, *hws-35*, and the *hws/hst* double mutant were pretreated with Mock or PIP, and systemic growth of *Psm* was assessed. As previously observed (Fig. 1), *hst-15* plants failed to induce systemic defenses (Fig. 4E, S4), whereas *hws-35* displayed WT-like PIP-induced systemic defenses (Fig. 4E, S4). Notably, the *hws/hst* double mutant also recovered WT-like PIP-induced defenses (Fig. 4C, S4).

miR159b, miR163, miR414, and miR822 are barely or not mis-regulated in *hst-15* mutants (Fig. S6), yet their mobility is enhanced in WT plants when defenses are induced (Fig. 3B, Fig. 3C). Thus, we then evaluated whether these miRNAs are required for a full systemic resistance response. To this end, we used T-DNA insertion mutants for *MIR163* and *MIR822*, as well as target mimicry transgenic lines for both miRNAs, which inhibit their activity and reduce their abundance (Todesco et al., 2010). Because their pronounced macroscopic phenotype prevented pathogen inoculation comparable to wild-type, miR159b mutants and mimicry lines were excluded from the analysis (Fig. S7A). As shown in Fig. 4F and S7B–D, mutating miR163 or miR822, or blocking their activity, reduced PIP-induced systemic defense. These results show that the activity of these miRNAs is required for systemic defense. They also suggest that in *hst-15* mutants the impaired defense results from their lack of movement.

### HST is enriched in the nucleus of PIP-treated plants

HST is localized in both the cytoplasm and nucleus, and its localization depends on the small GTPase RAN1 and the importin IMPA2 (Cambiagno *et al*., 2021). The role of HST in miRNA biogenesis and non-cell autonomous movement has been linked to its nuclear localization (Brioudes *et al*., 2021; Cambiagno *et al*., 2021; Gonzalo *et al*., 2022). To evaluate whether its subcellular localization affects the activation of systemic defenses, we assessed pathogen growth in WT, *ran1*, and *impa2* mutant plants previously treated with mock or PIP. Both *ran1* and *impa2* mutants showed a reduced induction of PIP-dependent systemic defenses compared to WT plants (Fig. 5A). Next, to assess whether impaired defenses in *ran1* and *impa2* result from altered HST localization, we analyzed HST subcellular distribution in PIP-treated tissues. Notably, HST was enriched in the nucleus to a similar extent as in *ran1* mutants (Fig. 5B), suggesting that its subcellular localization is modulated during the induction of systemic defenses.

**Figure 5:**
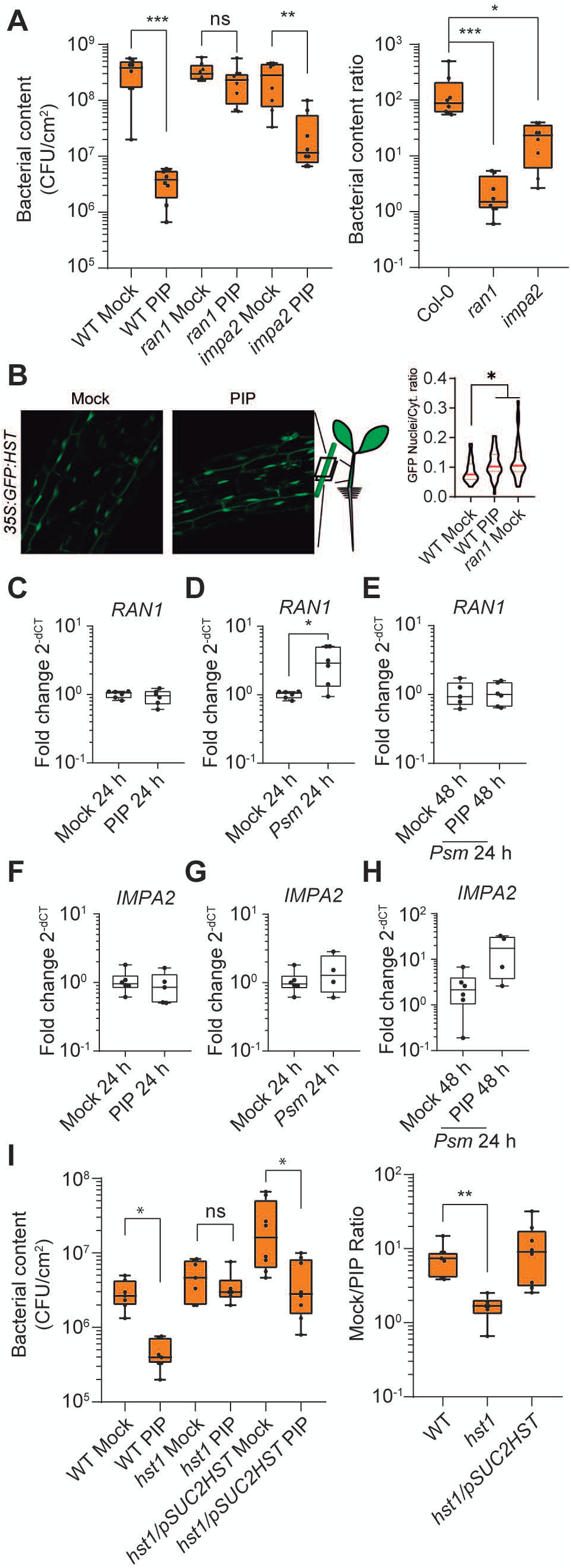
Subcellular localization of HST during systemic defense activation and its role in phloem companion cells. (A) *Pma* growth in distal leaves of wild-type (WT), *ran1*, and *impa2* plants plants pre-treated systemically with mock or PIP before pathogen inoculation. Pathogen growth is shown as CFU/cm^2^ (left) and as the ratio between Mock-and PIP-treated plants (right). (B) Subcellular localization of HST in seedlings grown on MS agar supplemented or not with PIP. Violin plot shows the ratio of GFP fluorescence intensity between nucleus and cytoplasm. HST localization in the *ran1* background was used as a control for nuclear enrichment. (C-H) Transcript levels of RAN1 and IMPA2 in vascular-enriched tissues from Mock-or PIP-treated plants (C, F), Psm-infected plants (D, G), or plants treated with Psm after Mock or PIP priming (E, H). (I) *Pma* growth in distal leaves of WT, *hst1*, and *hst1* plants complemented with *pSUC2:HST (hst1/pSUC2:HS7)*, as described in (A).*: p < 0.05, **: p < 0.01, ***: p < 0.001 by ANOVA (A, B) or Kruskal-Wallis test (A, I) with multiple comparisons corrected using Dunn’s test; or by two-tailed unpaired t-test (C-H).

A recent single-cell study reported RAN1 as a marker of phloem companion cells in infected leaves, while IMPA2 marks infected mesophyll cells [from supplementary data published (Delannoy et al., 2023; Zhu et al., 2023)]. Thus, we measured *RAN1* and *IMPA2* mRNA levels in vascular-enriched tissues from untreated leaves, systemically PIP-treated leaves, *Pma*-infected leaves, and leaves receiving both treatments. We confirmed vascular enrichment by comparing SUC2 (vascular) and ATML1 (epidermal) expression (Fig. S8). While in the vasculature of leaves from plants systemically treated with PIP did not induce *RAN1* or *IMPA2* expression (Fig. 5C, Fig. 5F), *Pma*-infected leaves increased *RAN1*-but not *IMPA2*-transcripts, consistent with previous findings (Delannoy *et al*., 2023) (Fig. 5D, Fig. 5G). Interestingly, pre-treatment with systemic PIP abolished the *RAN1* induction upon infection, while *IMPA2* transcripts showed a non-significant increase (Fig. 5E, Fig. 5H). These results suggest that SAR-inducing infection promotes *RAN1* transcript accumulation in companion cells, which is prevented by prior PIP-treatment. Conversely, *IMPA2* expression increases only in PIP-pretreated infected tissues. This highlights the importance of pathogen-driven manipulation of the RAN1/IMPA2 balance and HST localization in phloem companion cells. Future experiments will clarify differences between pathogen-and PIP-induced effects.

Since HST is a cell-autonomous protein (Brioudes *et al*., 2021), we tested whether its activity in CCs alone can trigger systemic defenses. We measured PIP-dependent systemic defense induction in *hst-1* mutants and in *hst-1* plants expressing HST exclusively in CCs (*hst-1/pSUC2:HST:GFP*). Notably, CC-specific expression of HST fully restored systemic defenses to WT levels (Fig. 5I). These results strongly suggest that HST’s within CCs is essential for systemic defense activation, where particularly its nuclear-cytoplasmic shuttling could be relevant.

## Discussion

The role of several miRNAs in plant defenses is increasingly evident, but the broader impact of miRNA biogenesis under stress remains understudied (Asadi and Millar, 2024). While immune defects in biogenesis mutants are often linked to reduced levels of specific miRNAs, dissecting global effects independent of individual miRNA abundance remains challenging. Here, we show that Arabidopsis systemic defenses are impaired in *hst* mutants, likely due to the loss of miRNA cell-to-cell movement. Our findings indicate that both the HST-dependent co-transcriptional processing of miRNA precursors and the distal movement of miRNAs are modulated in response to systemic defense induction in wild-type plants. Furthermore, HST subcellular localization is dynamically regulated during systemic defense activation, probably modulating its role in phloem CCs (Fig. 6).

**Figure 6:**
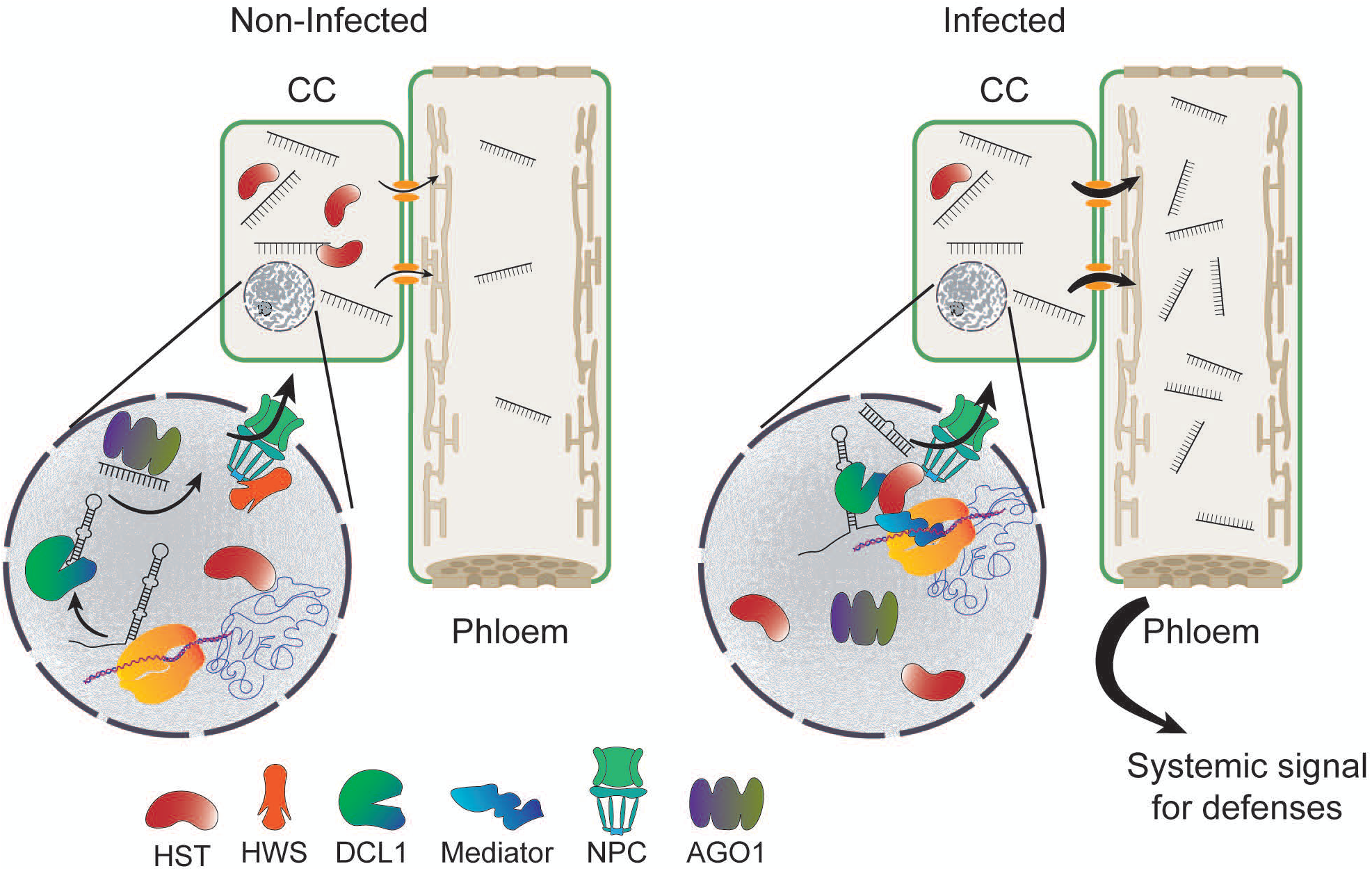
Model illustrating HST-dependent and-independent miRNA processing in Arabidopsis. In non-infected leaves, several primary miRNAs (pri-miRNAs) undergo post-transcriptional processing, likely independent of the nuclear pore complex (NPC), generating mature miRNAs with cell-autonomous activity. Upon infection, HST accumulates in the nucleus and promotes co-transcriptional processing of a subset of pri-miRNAs-including miR159, miR163, and miR822-probably at the NPC, thereby facilitating their phloem loading and long-distance movement. This HST-dependent pathway likely enables the systemic activation of plant defenses.

Apart from its role in systemic defenses, we found that HST also represses local defenses under mechanical stress. Among the miRNAs misregulated in *hst-15* (Cambiagno *et al*., 2021), few —including miR164b, miR167, miR169, and miR403—are known to influence plant defense by regulating their targets (NAC4, ARF6/8, NF-YA, and AGO2), which are upregulated in the mutant. Although these changes may contribute to local defense activation (Caruana et al., 2020; Hanemian et al., 2016; Harvey et al., 2011; Lee et al., 2017; Zhang et al., 2011b), they do not fully explain the impairment of systemic responses observed in *hst-15*. Moreover, when pathogen infection is initiated by spray inoculation, which mimics natural stomatal entry, mutants behave as WT plants. We also found no correlation between the differentially expressed miRNAs in *hst-15* and those in infected wild-type plants; thus, altered miRNA abundance alone cannot explain this phenotype, which is more likely caused by defective systemic miRNA movement. Supporting this, both the artificial miRNA amiRSUL and four endogenous miRNAs— miR159, miR163, miR414, and miR822—show increased phloem loading, suggesting enhanced distal mobility during defense induction. The phloem accumulation of most of these miRNAs was detected within the first 24 hours of systemic defense activation. Thus, they probably function as early mobile signals. This response appears selective, as miR171, miR402, and miR390 did not accumulate in phloem exudates following defense induction, suggesting that the movement of specific miRNA subsets contributes to the establishment or reinforcement of systemic immunity. Supporting this, it was recently shown that other sRNAs—such as miR390-derived tasiRNA-D7 and D8, and a 31-nt tRNA fragment (tRF31Asp2)—are specifically moved to distal leaves during SAR (Pujara, 2024; Shine *et al*., 2022). Similarly, the systemic movement of miR2111 from shoot to root improves nodulation in legumes (Tsikou *et al*., 2018). Together, these observations highlight the importance of mechanisms regulating sRNA distal movement in plant–microbe interactions. Our findings further suggest a general process that governs miRNA mobility and influences specific miRNA populations driving systemic plant defense.

Consistent with the induced mobility of miRNAs, we found that the precursors of miR159, miR163, and miR822—but not miR171—are preferentially co-transcriptionally processed during PIP-induced defense activation. This agrees with previous reports showing that cold or heat stress shifts the balance between co-and post-transcriptional processing, thereby influencing their non-cell-autonomous activity (Gonzalo *et al*., 2025; Gonzalo *et al*., 2022). Beyond HST’s role in pri-miRNA processing at the nuclear pore complex (NPC) and the influence of microtubule dynamics on AGO1 loading (Brioudes *et al*., 2021; Devers *et al*., 2020; Fan *et al*., 2022; Gonzalo *et al*., 2025), the mechanisms underlying the non-cell-autonomous activity of miRNAs are still largely unknown. Even less is known about how miRNAs are selectively mobilized across tissues, developmental stages, or in response to stresses (Loreti and Perata, 2022; Voinnet, 2022). Our findings suggest a nuclear export fine-tuning mechanism that controls the movement of specific miRNAs involved in pathogen resistance and may operate differently under other stress conditions. The systemic movement miR159b, miR163, miR414, and miR822 was found to be HST-dependent. Moreover, analysis of the *hws35/hst-15* double mutant—in which miRNA mobility is restored while the miRNA abundance characteristic of *hst-15* remains unchanged (Gonzalo *et al*., 2025)—revealed that systemic defense induction is also recovered. Together, these results strongly suggest that HST is a key component in the movement of a subset of mobile miRNAs important for systemic defense, while additional regulatory factors may further refine which specific miRNAs are mobilized. In contrast, other small RNAs known to move systemically—such as tasiRNA-D7 and D8, and tRNA-derived fragments involved in SAR (Pujara, 2024; Shine *et al*., 2022)—likely rely on distinct pathways, as their movement is independent of HST (Brioudes *et al*., 2021). Therefore, the systemic defense defects observed in *hst* mutants are most likely due to impaired miRNA movement rather than defects in the trafficking of other sRNA types.

miRNA movement involves symplastic transport through plasmodesmata and long-distance spreading via the phloem. For phloem loading in Arabidopsis, miRNAs from parenchyma cells must reach CCs, but the scarcity of plasmodesmata between these cell types suggests that transfer may occur through apoplastic EVs (Voinnet, 2025). This route likely entails initial symplastic movement within parenchyma, followed by EV-mediated export to the apoplast and final uptake into CCs, although the regulatory mechanisms remain poorly understood (Voinnet, 2025). In this context, HST may facilitate miRNA symplastic movement either within parenchyma or between CCs and the phloem. Supporting this, we found that HST expression specifically in phloem CCs is sufficient to restore systemic defenses. Additional experiments expressing HST in cell types other than CCs would help clarify this mechanism. Altogether, these findings suggest that the induction of systemic defenses involves multiple mechanisms that coordinate the movement of different sRNA classes, likely with overlapping functions to reinforce systemic programs. Uncovering the potential roles of other components beyond HST, like specific AGOs or plasmodesmata-associated components, will help clarify the fine-tuned regulation of sRNA movement as a signaling mechanism in plant stress responses.

Along with the increased HST-dependent mobility of miR163 and miR822, our study demonstrates that both are required for the induction of systemic resistance. The roles of miR822 and miR163 in plant–pathogen interactions remain largely unexplored, as do the functions of their mRNA targets. miR822 follows the canonical miRNA biogenesis pathway, although uniquely in a DCL4-dependent manner (Zhang et al., 2010). It is loaded onto AGO9 and AGO1 (Fan *et al*., 2022; Li et al., 2010; Tovar-Aguilar et al., 2024), and shows mild induction during non-host interactions, pattern-triggered immunity (PTI), and effector-triggered immunity (ETI) responses (Zhang *et al*., 2010). Interestingly, miR822 association with AGO1 increases upon flg22 treatment (Li *et al*., 2010), potentially modulating both its activity and systemic mobility. miR163, a 24-nt miRNA preferentially loaded onto AGO1 and AGO2 (Jeong et al., 2013), is induced by *Pst* infection and salicylic acid (SA) treatment. It has been proposed to act as a negative regulator of immunity, potentially fine-tuning defense outputs (Chow and Ng, 2017; Litholdo et al., 2018). The observation of altered AGO loading in infected tissues (Fan *et al*., 2022; Li *et al*., 2010; Tovar-Aguilar *et al*., 2024), along with the HST-dependent mobility demonstrated here, points to a functional role for miR822 and miR163 in long-distance immune signaling. Furthermore, we speculate that these miRNAs—and potentially others—may selectively load into different AGOs under stress conditions, providing insight into the mechanisms that regulate the mobility of specific miRNA subsets.

We show that the increased HST nuclear localization in PIP-treated plants correlates with enhanced miRNA mobility and defense induction. Accordingly, *ran1* and *impa2* mutants, which alter HST nucleo-cytoplasmic localization, are also essential for systemic resistance. In support, both RAN1 and IMPA2 have been implicated in defense induction. RAN1 and IMPA2 contribute to resistance against *Rhizoctonia solani* (Parween and Sahu, 2022), with RAN1 also playing a role in defense against *Pseudomonas syringae pv. tomato* (Xu et al., 2021). Additionally, IMPA1, IMPA2, and IMPA4 act redundantly to suppress defense-related gene expression and resistance to *Colletotrichum higginsianum* (Mori et al., 2024). In contrast, studies using the *hst-3* allele suggested that HST function is independent of its nuclear or cytoplasmic localization (Brioudes *et al*., 2021). However, the *hst-3* allele is defective in HST–RAN1 interaction while maintaining normal subcellular distribution in roots of non-infected plants. Thus, one possibility is that RAN1-and IMPA2-dependent regulation of HST subcellular localization could modulate its role in miRNA cell-to-cell movement, a mechanism that may be particularly relevant under stress conditions. Supporting this, previous studies reported RAN1 transcript accumulation in phloem companion cells of infected plants (Delannoy *et al*., 2023; Zhu *et al*., 2023). In line with this, we found that *Psm*, PIP, and their combination modulate RAN1 and IMPA2 transcript levels in vascular tissues, reinforcing a central role of HST subcellular localization. Even when *hst-15* plants expressing HST at CCs restored systemic defenses, its nuclear accumulation extended beyond CCs, consistent with IMPA2 transcript presence in infected mesophyll cells (Zhu *et al*., 2023). This suggests that RAN1-and IMPA2-dependent regulation of HST localization supports nucleus– cytoplasm shuttling in multiple tissues and particularly in CCs, where HST likely facilitates miRNA loading into the phloem for systemic defense.

Overall, our results show that HST is a positive regulator of systemic responses, mediating the non-cell-autonomous activity of specific miRNAs by facilitating their loading into the phloem, ultimately enabling their distal movement. Future studies aimed at deciphering how global mechanisms involved in miRNA biogenesis and mobility are fine-tuned to allow the long-distance movement of selected miRNAs will help clarify the role of miRNAs in the fine regulation of plant immunity.

## Materials and Methods

### Plant materials and growth conditions

Seeds of *Arabidopsis thaliana* Col-0 (CS22681), *hst-15* (SALK_079290), *hst-3* (CS24278), *hst-1* (CS3810), *ran1-1* (SALK_138680), *ran1-2* (SALK_067649), *impa2* (SALK_099707), *mir163* (SALK_034556C), *mir822* (SALK_023928), *hws-35* (EMS mutation), and transgenic plants harboring *pSUC2:amiRSUL*, *35S:MIM163*, *35S:MIM822*, *35S:GFP:HST*, *pSUC2:HST:GFP* (Table S1) were used in this study. Seeds were sterilized with 10% (v/v) aqueous bleach and 0.5% SDS, and stratified for 2–3 days at 4 °C. Germination was performed on plates containing 2.2 g/l Murashige– Skoog (MS) medium (pH 5.7) with 0.6% agar for seedling assays, or in soil (Growmix substrate) for four-week-old plant assays.

### Pathogen and chemical treatments, and systemic resistance assays

Four-week-old plants grown in pots were treated by soil flooding with 1 mM PIP or water (control) for 30 minutes, avoiding contact with aerial tissues (Miranda de la Torre et al., 2023). Pots were then drained and returned to the growth chamber. One day later, leaves were syringe-infiltrated with 5 x 10^5^ CFU/ml of *Pseudomonas cannabina* pv. *alisalensis* (*PmaDG3*; formerly *P. syringae* pv. *maculicola* (Baltrus *et al*., 2011)). Only when highligjhted (Fig. E, F). pathogen was spray inoculated at 1 x 10^8^ CFU/ml. After three days, bacterial content from three leaves per sample (six leaf discs, eight samples per experiment) was quantified by extraction in 10 mM MgCl_2_, followed by plating of serial dilutions on King’s B medium supplemented with rifampicin. For SAR assays, instead of PIP treatment, *Pseudomonas syringae* pv. *tomato* harboring the avirulence factor AvrRpt2 (*Pst-AvrRpt2*) was infiltrated into lower leaves at 5 x 10^6^ CFU/ml. Forty-eight hours post-infection, upper leaves were infiltrated with *Pma* to assess pathogen growth three days later. The ratio of pathogen growth between mock and PIP-treated or SAR-induced plants was calculated as the mean CFU in the mock condition divided by the CFU in each PIP-treated or SAR-induced sample. Statistical significance (*p < 0.05; **p < 0.01; ***p < 0.001) was evaluated using ANOVA or Kruskal–Wallis tests (depending on Shapiro–Wilk normality test), with multiple comparisons corrected using Dunn’s test. T-tests or Mann–Whitney tests were used for pairwise comparisons.

When defense induction by *Pma*, *Pst-AvrRpt2*, or PIP was tested in infiltrated leaves, the same concentrations were used for syringe infiltration.

### RNA-seq analysis

Previously published mRNA-seq data from Cambiagno et al. (Cambiagno *et al*., 2021) were used for GO-term analysis. A PANTHER Overrepresentation Test (Slim Biological Process) was applied to the upregulated genes in *hst-15* (DESeq2, log2 fold change > 0.6; adjusted p-value < 0.05). Fold enrichment and FDR values were visualized using dot plots in RStudio. Additional RNA-seq analyses were performed using raw data from ERP127706 (Yildiz *et al*., 2021), ERP105389 (Hartmann *et al*., 2018), SRP217896 (Desrut *et al*., 2020), SRP258528, ERP013651 (Bernsdorff *et al*., 2016), ERP115669 (Baum *et al*., 2019), and SRP123046 (Stringlis *et al*., 2018). Reads were mapped to the *A. thaliana* reference genome (TAIR10) using STAR (Dobin et al., 2013), and gene counts were obtained using featureCounts. All MIR genes annotated in the GFF3 files were included in the analysis. Differential gene expression was determined with DESeq2 in RStudio. MIR genes with adjusted p-value < 0.05 were considered in subsequent analyses. Common up-or downregulated MIR genes across datasets were visualized with UpSet plots in RStudio. For Fig. 1H, pri-miRNAs differentially expressed in mRNA-seq of systemically treated leaves were plotted with RStudio and used as indirect indicators of miRNA abundance. The twelve mRNA-seq datasets from systemically defense-induced tissues (Baum *et al*., 2019; Bernsdorff *et al*., 2016; Desrut *et al*., 2020; Hartmann *et al*., 2018; Yildiz *et al*., 2021), were compared against miRNAs mis-regulated in *hst-15* (Cambiagno *et al*., 2021). MIR genes downregulated in systemic defense-induced tissues (from forty mis-regulated) corresponded to eight miRNAs downregulated in *hst-15* (highlighted dots, Fig. 1H). In Fig. S1, the abundance of the genes targeted by miR164b, miR167, miR169, and miR403 (previously associated with plant defenses) was analyzed from *hst-15* mRNA-seq and plotted with RStudio.

sRNA-seq data from *hst-15*, *hws35/hst-15* (Gonzalo *et al*., 2025) (PRJEB90908) and WT plants, either infected or mock-treated with *PstAvrRpt2* at 14 h post-infection (Zhang *et al*., 2011a) (PRJNA122227), were analyzed using Galaxy (Galaxy, 2024). Reads were mapped to the TAIR10 genome with miRDeep2 Mapper and quantified using miRDeep2 Quantifier, with Arabidopsis pri-miRNA and mature miRNA sequences from miRBase (release 22.1) as input. Differential miRNA abundance was assessed with DESeq2 (log2 fold change > 1 or < −1; adjusted p < 0.05). In Fig. 4D, pathogen-responsive genes refer to those from PRJNA122227 (Zhang *et al*., 2011a) that, in our analysis, show p < 0.05

### Quantification of leaf bleaching in pSUC2:amiRSUL plants

Leaf bleaching around veins in transgenic *pSUC2:amiRSUL* plants was quantified as previously described (Fan *et al*., 2022), with modifications. Images of individual leaves were captured using an Epson CX5600 scanner and processed in Fiji/ImageJ. For total leaf area, leaves were segmented from the background using the Color Threshold plugin, followed by Clear Outside, removal of small particles with Analyze Particles, and erosion using the EDM Erosion plugin from BioVoxxel (10 iterations). The Close function was applied to complete the leaf contour if needed. To segment bleached areas, the leaf mask was reapplied to the original image to remove margins using Clear Outside. The image was converted to 8-bit grayscale, gamma-corrected (2.75), followed by Subtract Background (rolling ball radius = 100). Bleached regions were segmented using the Isodata algorithm. Binary bleached masks were generated, and filters were applied to remove artifacts such as trichomesand. The ratio of bleached area to total leaf area was plotted. Statistical differences (*p < 0.05; **p < 0.01 or ***p < 0.001) were calculated using ANOVA and corrected for multiple comparisons using Dunńs test. Leaves with bleached/total area similar to the media of all the replicates of a treatment are shown. Yellow color intensity in representative leaf veins was quantified using ImageJ. A line was manually drawn along the same corresponding vein on each leaf across treatments, and the resulting intensity profile was extracted with the Plot Profile function. Yellow intensity was calculated as the mean of the red and green channels using the formula (red + green)/2.

### RNA analysis

RNA was extracted using TRIzol reagent. RT-qPCR for miRNAs was performed using stem-loop RT primers (Varkonyi-Gasic et al., 2007). Briefly, cDNA was synthesized from 1 µg of total RNA using the RevertAid RT Reverse Transcription Kit (Thermo Fisher Scientific), with specific stem-loop primers and oligo-dT included in the same reaction (Table S2). qPCRs were performed with at least three biological replicates, using as an internal control *Actin2/8* for cDNA from total leaves or UBC9 for cDNA from phloem exudates. Fold changes were calculated using the 2^-ΔΔCt^ method and presented as box plots. Statistical differences (*p < 0.05; **p < 0.01 or ***p < 0.001) were calculated using ANOVA or Krusal-Wallis (depending Shapiro-Wilk normality test) and corrected for multiple comparisons using Dunńs test. Two-tailed unpaired *t*-test or Mann-Whitney tests were used for comparing pairs of samples.

### Phloem exudate collection

Phloem exudates were collected using the EDTA-facilitated method (Tetyuk et al., 2013). Briefly, 16 leaves from one-month-old plants were sliced with a razor blade and preincubated in Petri dishes containing 20 mM K₂-EDTA. Eight leaves were then transferred to 1.5 ml tubes with 20 mM K₂-EDTA (two tubes per sample) for 1 h in a humid chamber under regular temperature and light conditions. Leaves were washed with Milli-Q water and incubated in new tubes for exudates collection containing water and Riboblock RNase inhibitor (Thermo) for 24 h in a humid chamber. Samples treated with mock (MEX), pathogen (PEX), or PIP (PIPEX) were used for RNA extraction with TRIzol.

### Vascular tissue separation from leaves

To enrich vascular tissue, the Meselect protocol was used (Svozil et al., 2016). Leaves from one-month-old plants were placed between paper tapes to separate the lower epidermis. Vascular tissue was enriched using Cellulase and Macerozyme (Onozuca) treatments to digest mesophyll tissue. Tapes with lower epidermal and vascular tissues were processed separately for RNA extraction and qPCR.

### Chromatin-bound and nucleoplasmic pri-miRNA extraction

To analyze co-transcriptional processing, chromatin-bound and nucleoplasmic pri-miRNAs were extracted as described (Zhang et al., 2022). Nuclear-enriched samples were washed with 1 M urea to remove unbound RNAs and proteins. Chromatin-bound RNAs (pellet) and unbound RNAs (supernatant) were subjected to RNA extraction and qPCR. The abundance of processed versus unprocessed pri-miRNAs was inferred as described (Gonzalo *et al*., 2022). Briefly, RT-qPCR was performed using primers targeting the stem-loop region and, in parallel, the region surrounding the DCL cleavage site. *Actin2/8* was used as an internal control.

### Microscopy

Subcellular localization of HST in mock-, *Pma*-, or PIP-treated plants was analyzed as previously described (Cambiagno *et al*., 2021). GFP was excited at 480 nm, and emission was collected at 520/30 nm using Nikon Eclypse CSI confocal microscopy. GFP fluorescence intensity was measured in the nucleus and cytoplasm of ∼45 hypocotyl cells from ten plants. Mean signal intensities were quantified using ImageJ, and nucleus-to-cytoplasm ratios were plotted. Statistical differences (*p < 0.05; **p < 0.01 or ***p < 0.001) were calculated using ANOVA and corrected for multiple comparisons using Dunńs test.

**Figure S1: miRNAs and defense-related genes misregulated in *hst-15* and in samples with systemically induced defenses.** Box plot showing normalized counts of genes targeted by the six miRNAs labeled in Fig. 1H, in non-treated *hst-15* plants. Significant differences are based on DESeq2 analysis.

**Figure S2: Selection of miRNAs involved in local and systemic defense activation.**

(A) Upset plot showing *pri*-miRNAs that are upregulated (top) or downregulated (bottom) in samples treated locally or systemically with different defense inducers. *MIR* genes misregulated in more than nine transcriptomes are highlighted. (B) miRNA abundance in leaves untreated (-), treated systemically with PIP for 24 h, locally infected with *Pma* for 24 or 48 h, or co-treated with PIP and *Pma* (48 h with PIP and by 24 h with *Pma*, or 96 h with PIP and 72 h with *Pma*). (C) miRNA abundance measured by RT-qPCR in phloem exudates collected from Mock-treated (MEX), or PIP-treated (PIPEX) leaves. *: p < 0.05, **: p < 0.01, ***: p < 0.001 by ANOVA with multiple comparisons corrected using Dunn’s test (B) or by two-tailed unpaired t-test (C).

**Figure S3: Detection of miR171 and Actin in phloem exudates.**

Abundance of miR171 and Actin in phloem exudates collected from Mock-treated (MEX) or PIP-treated (PIPEX) leaves of wild-type (WT) and *hst-15* plants. These are negative controls for Figure 5B. *: p < 0.05 by ANOVA with multiple comparisons corrected using Dunn’s test.

**Figure S4: Pathogen growth in *hst-15* plants complemented with the *hws-35* mutation.**

*Pma* growth in leaves of wild-type (WT), *hst-15*, and *hws-35/hst-15* (*hws/hst*) plants plants pre-treated systemically with mock or PIP before pathogen inoculation. Pathogen growth is expressed as CFU/cm². Statistical significance: *: p < 0.05, **: p < 0.01, determined by Mann–Whitney test.

**Figure S5: Restoration of miRNA phloem loading in *hws35/hst-15* double mutant.**

Abundance of miRNAs in phloem exudates collected from Mock-treated (MEX) or PIP-treated (PIPEX) leaves of wild-type (WT), *hst-15*, and *hws-35/hst-15* (*hws/hst*) double mutant plants. Fold change was calculated comparing PIP-treated to Mock-treated samples (indicated by the red line) for each genotype. *: p < 0.05, **: p < 0.01, ***: p < 0.001 by Kruskal–Wallis test with multiple comparisons corrected using Dunn’s test.

**Figure S6. miRNA abundance in *hst-15*.**

Abundance of miR159b, miR163, miR414, and miR822 in *hst-15* plants from Cambiagno et al 2021 (*a), or Gonzalo et al., 2025 (*b). Fold change and False Discovery Rate (FDR) values are indicated above each bar.

**Figure S7. Pathogen growth in plants with impaired miR163 or miR822 activity.**

(A) Phenotype of 45-day-old wild-type and *MIM159b* plants. White bars represent 0.5 cm. (B) *Pma* growth in leaves of wild-type (WT), *MIM163*, *MIM822*, *mir163*, and *mir822* plants pre-treated systemically with mock or PIP before pathogen inoculation. Bacterial content is expressed as CFU/cm². Statistical significance: *: p < 0.05, **: p < 0.01, ***: p < 0.001, determined by ANOVA with multiple comparisons corrected using Dunn’s test.

**Figure S8. Enrichment of vasculature and epidermal cell.**

Transcript levels of *SUC2* and *ATML1* in vascular-and epidermal-enriched tissues. *: *p* < 0.05, **: *p* < 0.01 by ANOVA test with multiple comparisons corrected using Dunn’s test.

**Table S1: Wild-type and mutant plants used in this study.**

**Table S2: Primers used in this study.**

## Founding

This work was supported by grants from the Agencia Nacional de Promocion Cientifica y Tecnologica (PICT-2019-0095, PICT-2019-0533, PICT-2020-0071, PICT-2021-0286) and Instituto Nacional de Tecnologías Agropecuarias (2023-PDI083) to DAC, and Consejo Nacional de Investigaciones Científicas y Técnicas de Argentina (PIP-GI 11220200102984CO) to HRL. AT, NMC, HRL, and DAC are members of CONICET, MM, and LQ are fellows of CONICET, and NA is an alumni.

## Author contribution

MM, NA, and LQ performed most of the experiments. AT optimized the methodology and performed the quantification of leaf bleaching. NMC, HRL, and DAC conceived the study. MM, NMC, and DAC wrote the manuscript. HRL and DAC secure the founds, and DAC supervised the experimental procedures.

## Acknowledgement

We thank Leandro I. Ortega for assistance with microscopy and Virginia Lobatto for technical support. We are grateful to Pablo Manavella for critically reading the manuscript and, together with Olivier Voinnet, for providing seeds essential to the conception of this study. We also thank FC for valuable support in project development.

## Declaration of Interests

The authors declare no competing interests.

## Declaration of Generative AI and AI-assisted technologies in the writing process

During the preparation of this work the authors used Chat-GPT 4.0 to improve readability and language, and for improving the presentation of some plots in R. After using this tool/service, the authors reviewed and edited the content as needed and take full responsibility for the content of the publication.

## Notes

### Competing Interest Statement

The authors have declared no competing interest.

## References

Achkar, N.P., Cambiagno, D.A., and Manavella, P.A. (2016). miRNA Biogenesis: A Dynamic Pathway. Trends Plant Sci 21:1034–1044. 10.1016/j.tplants.2016.09.003.

Asadi, M., and Millar, A.A. (2024). Review: Plant microRNAs in pathogen defense: A panacea or a piece of the puzzle? Plant Sci 341:111993. 10.1016/j.plantsci.2024.111993.

Bakirbas, A., Castro-Rodriguez, R., and Walker, E.L. (2023). The Small RNA Component of Arabidopsis thaliana Phloem Sap and Its Response to Iron Deficiency. Plants (Basel) 1210.3390/plants12152782.

Baltrus, D.A., Nishimura, M.T., Romanchuk, A., Chang, J.H., Mukhtar, M.S., Cherkis, K., Roach, J., Grant, S.R., Jones, C.D., and Dangl, J.L. (2011). Dynamic evolution of pathogenicity revealed by sequencing and comparative genomics of 19 Pseudomonas syringae isolates. PLoS Pathog 7:e1002132. 10.1371/journal.ppat.1002132.

Baum, S., Reimer-Michalski, E.M., Bolger, A., Mantai, A.J., Benes, V., Usadel, B., and Conrath, U. (2019). Isolation of Open Chromatin Identifies Regulators of Systemic Acquired Resistance. Plant Physiol 181:817–833. 10.1104/pp.19.00673.

Bernsdorff, F., Doring, A.C., Gruner, K., Schuck, S., Brautigam, A., and Zeier, J. (2016). Pipecolic Acid Orchestrates Plant Systemic Acquired Resistance and Defense Priming via Salicylic Acid-Dependent and-Independent Pathways. Plant Cell 28:102–129. 10.1105/tpc.15.00496.

Brioudes, F., Jay, F., Sarazin, A., Grentzinger, T., Devers, E.A., and Voinnet, O. (2021). HASTY, the Arabidopsis EXPORTIN5 ortholog, regulates cell-to-cell and vascular microRNA movement. EMBO J 40:e107455. 10.15252/embj.2020107455.

Cai, Q., Halilovic, L., Shi, T., Chen, A., He, B., Wu, H., and Jin, H. (2023). Extracellular vesicles: cross-organismal RNA trafficking in plants, microbes, and mammalian cells. Extracell Vesicles Circ Nucl Acids 4:262–282. 10.20517/evcna.2023.10.

Cambiagno, D.A., Giudicatti, A.J., Arce, A.L., Gagliardi, D., Li, L., Yuan, W., Lundberg, D.S., Weigel, D., and Manavella, P.A. (2021). HASTY modulates miRNA biogenesis by linking pri-miRNA transcription and processing. Mol Plant 14:426–439. 10.1016/j.molp.2020.12.019.

Caruana, J.C., Dhar, N., and Raina, R. (2020). Overexpression of Arabidopsis microRNA167 induces salicylic acid-dependent defense against Pseudomonas syringae through the regulation of its targets ARF6 and ARF8. Plant Direct 4:e00270. 10.1002/pld3.270.

Chow, H.T., and Ng, D.W. (2017). Regulation of miR163 and its targets in defense against Pseudomonas syringae in Arabidopsis thaliana. Sci Rep 7:46433. 10.1038/srep46433.

Conrath, U., Beckers, G.J., Langenbach, C.J., and Jaskiewicz, M.R. (2015). Priming for enhanced defense. Annu Rev Phytopathol 53:97–119. 10.1146/annurev-phyto-080614-120132.

de Felippes, F.F., Ot, F., and Weigel, D. (2011). Comparative analysis of non-autonomous effects of tasiRNAs and miRNAs in Arabidopsis thaliana. Nucleic Acids Res 39:2880–2889. 10.1093/nar/gkq1240.

Delannoy, E., Batardiere, B., Pateyron, S., Soubigou-Taconnat, L., Chiquet, J., Colcombet, J., and Lang, J. (2023). Cell specialization and coordination in Arabidopsis leaves upon pathogenic attack revealed by scRNA-seq. Plant Commun 4:100676. 10.1016/j.xplc.2023.100676.

Desrut, A., Moumen, B., Thibault, F., Le Hir, R., Coutos-Thevenot, P., and Vriet, C. (2020). Beneficial rhizobacteria Pseudomonas simiae WCS417 induce major transcriptional changes in plant sugar transport. J Exp Bot 71:7301–7315. 10.1093/jxb/eraa396.

Devers, E.A., Brosnan, C.A., Sarazin, A., Albertini, D., Amsler, A.C., Brioudes, F., Jullien, P.E., Lim, P., Schott, G., and Voinnet, O. (2020). Movement and differential consumption of short interfering RNA duplexes underlie mobile RNA interference. Nat Plants 6:789–799. 10.1038/s41477-020-0687-2.

Dobin, A., Davis, C.A., Schlesinger, F., Drenkow, J., Zaleski, C., Jha, S., Batut, P., Chaisson, M., and Gingeras, T.R. (2013). STAR: ultrafast universal RNA-seq aligner. Bioinformatics 29:15–21. 10.1093/bioinformatics/bts635.

Fan, L., Zhang, C., Gao, B., Zhang, Y., Stewart, E., Jez, J., Nakajima, K., and Chen, X. (2022). Microtubules promote the non-cell autonomous action of microRNAs by inhibiting their cytoplasmic loading onto ARGONAUTE1 in Arabidopsis. Dev Cell 57:995–1008 e1005. 10.1016/j.devcel.2022.03.015.

Fang, X., Cui, Y., Li, Y., and Qi, Y. (2015). Transcription and processing of primary microRNAs are coupled by Elongator complex in Arabidopsis. Nat Plants 1:15075. 10.1038/nplants.2015.75.

Foret, J., Kim, J.G., Sattely, E.S., and Mudgett, M.B. (2024). Transcriptome analysis reveals role of transcription factor WRKY70 in early N-hydroxy-pipecolic acid signaling. Plant Physiol 19710.1093/plphys/kiae544.

Galaxy, C. (2024). The Galaxy platiorm for accessible, reproducible, and collaborative data analyses: 2024 update. Nucleic Acids Res 52:W83–W94. 10.1093/nar/gkae410.

Gonzalo, L., Gagliardi, D., Zlauvinen, C., Gulanicz, T., Arce, A.L., Fernandez, J., Cambiagno, D.A., Merchante, C., Zienkiewicz, A., Jarmolowski, A., et al. (2025). The nuclear pore complex acts as a hub for pri-miRNA transcription and processing in plants. Nucleic Acids Res 5310.1093/nar/gkaf885.

Gonzalo, L., Tossolini, I., Gulanicz, T., Cambiagno, D.A., Kasprowicz-Maluski, A., Smolinski, D.J., Mammarella, M.F., Ariel, F.D., Marquardt, S., Szweykowska-Kulinska, Z., et al. (2022). R-loops at microRNA encoding loci promote co-transcriptional processing of pri-miRNAs in plants. Nat Plants 8:402–418. 10.1038/s41477-022-01125-x.

Hanemian, M., Barlet, X., Sorin, C., Yadeta, K.A., Keller, H., Favery, B., Simon, R., Thomma, B.P., Hartmann, C., Crespi, M., et al. (2016). Arabidopsis CLAVATA1 and CLAVATA2 receptors contribute to Ralstonia solanacearum pathogenicity through a miR169-dependent pathway. New Phytol 211:502–515. 10.1111/nph.13913.

Hartmann, M., Zeier, T., Bernsdorff, F., Reichel-Deland, V., Kim, D., Hohmann, M., Scholten, N., Schuck, S., Brautigam, A., Holzel, T., et al. (2018). Flavin Monooxygenase-Generated N-Hydroxypipecolic Acid Is a Critical Element of Plant Systemic Immunity. Cell 173:456–469 e416. 10.1016/j.cell.2018.02.049.

Harvey, J.J., Lewsey, M.G., Patel, K., Westwood, J., Heimstadt, S., Carr, J.P., and Baulcombe, D.C. (2011). An antiviral defense role of AGO2 in plants. PLoS One 6:e14639. 10.1371/journal.pone.0014639.

Jeong, D.H., Thatcher, S.R., Brown, R.S., Zhai, J., Park, S., Rymarquis, L.A., Meyers, B.C., and Green, P.J. (2013). Comprehensive investigation of microRNAs enhanced by analysis of sequence variants, expression patterns, ARGONAUTE loading, and target cleavage. Plant Physiol 162:1225–1245. 10.1104/pp.113.219873.

Jones, J.D., Vance, R.E., and Dangl, J.L. (2016). Intracellular innate immune surveillance devices in plants and animals. Science 35410.1126/science.aaf6395.

Lee, M.H., Jeon, H.S., Kim, H.G., and Park, O.K. (2017). An Arabidopsis NAC transcription factor NAC4 promotes pathogen-induced cell death under negative regulation by microRNA164. New Phytol 214:343–360. 10.1111/nph.14371.

Li, S., Wang, X., Xu, W., Liu, T., Cai, C., Chen, L., Clark, C.B., and Ma, J. (2021). Unidirectional movement of small RNAs from shoots to roots in interspecific heterogratis. Nat Plants 7:50–59. 10.1038/s41477-020-00829-2.

Li, Y., Zhang, Q., Zhang, J., Wu, L., Qi, Y., and Zhou, J.M. (2010). Identification of microRNAs involved in pathogen-associated molecular pattern-triggered plant innate immunity. Plant Physiol 152:2222–2231. 10.1104/pp.109.151803.

Litholdo, C.G., Jr., Eamens, A.L., and Waterhouse, P.M. (2018). The phenotypic and molecular assessment of the non-conserved Arabidopsis MICRORNA163/S-ADENOSYL-METHYLTRANSFERASE regulatory module during biotic stress. Mol Genet Genomics 293:503–523. 10.1007/s00438-017-1399-9.

Loreti, E., and Perata, P. (2022). Mobile plant microRNAs allow communication within and between organisms. New Phytol 235:2176–2182. 10.1111/nph.18360.

Macho, A.P., and Zipfel, C. (2014). Plant PRRs and the activation of innate immune signaling. Mol Cell 54:263–272. 10.1016/j.molcel.2014.03.028.

Miranda de la Torre, J.O., Peppino Margutti, M.Y., Lescano Lopez, I., Cambiagno, D.A., Alvarez, M.E., and Cecchini, N.M. (2023). The Arabidopsis chromatin regulator MOM1 is a negative component of the defense priming induced by AZA, BABA and PIP. Front Plant Sci 14:1133327. 10.3389/fpls.2023.1133327.

Mori, A., Nakagawa, S., Suzuki, T., Suzuki, T., Gaudin, V., Matsuura, T., Ikeda, Y., and Tamura, K. (2024). The importin alpha proteins IMPA1, IMPA2, and IMPA4 play redundant roles in suppressing autoimmunity in Arabidopsis thaliana. Plant J 10.1111/tpj.17203.

Park, M.Y., Wu, G., Gonzalez-Sulser, A., Vaucheret, H., and Poethig, R.S. (2005). Nuclear processing and export of microRNAs in Arabidopsis. Proc Natl Acad Sci U S A 102:3691–3696. 10.1073/pnas.0405570102.

Parperides, E., El Mounadi, K., and Garcia-Ruiz, H. (2023). Induction and suppression of gene silencing in plants by nonviral microbes. Mol Plant Pathol 24:1347–1356. 10.1111/mpp.13362.

Parween, D., and Sahu, B.B. (2022). An Arabidopsis nonhost resistance gene, IMPORTIN ALPHA 2 provides immunity against rice sheath blight pathogen, Rhizoctonia solani. Curr Res Microb Sci 3:100109. 10.1016/j.crmicr.2022.100109.

Pelaez, P., and Sanchez, F. (2013). Small RNAs in plant defense responses during viral and bacterial interactions: similarities and differences. Front Plant Sci 4:343. 10.3389/fpls.2013.00343.

Pujara, D. (2024). Nuclear tRNA-derived RNA fragments (tRFs) trigger immunity in arabidopsis and potentially function as a mobile signal in systemic acquired resistance. Doctoral dissertation, Texas State University.

Rebolledo-Prudencio, O.G., Estrada-Rivera, M., Dautt-Castro, M., Arteaga-Vazquez, M.A., Arenas-Huertero, C., Rosendo-Vargas, M.M., Jin, H., and Casas-Flores, S. (2021). The small RNA-mediated gene silencing machinery is required in Arabidopsis for stimulation of growth, systemic disease resistance, and suppression of the nitrile-specifier gene NSP4 by Trichoderma atroviride. Plant J 10.1111/tpj.15599.

Sarris, P.F., Trantas, E.A., Baltrus, D.A., Bull, C.T., Wechter, W.P., Yan, S., Ververidis, F., Almeida, N.F., Jones, C.D., Dangl, J.L., et al. (2013). Comparative genomics of multiple strains of Pseudomonas cannabina pv. alisalensis, a potential model pathogen of both monocots and dicots. PLoS One 8:e59366. 10.1371/journal.pone.0059366.

Shine, M.B., Zhang, K., Liu, H., Lim, G.H., Xia, F., Yu, K., Hunt, A.G., Kachroo, A., and Kachroo, P. (2022). Phased small RNA-mediated systemic signaling in plants. Sci Adv 8:eabm8791. 10.1126/sciadv.abm8791.

Silvestri, A., Bansal, C., and Rubio-Somoza, I. (2024). Atier silencing suppression: miRNA targets strike back. Trends Plant Sci 10.1016/j.tplants.2024.05.001.

Song, L., Fang, Y., Chen, L., Wang, J., and Chen, X. (2021). Role of non-coding RNAs in plant immunity. Plant Commun 2:100180. 10.1016/j.xplc.2021.100180.

Stringlis, I.A., Proietti, S., Hickman, R., Van Verk, M.C., Zamioudis, C., and Pieterse, C.M.J. (2018). Root transcriptional dynamics induced by beneficial rhizobacteria and microbial immune elicitors reveal signatures of adaptation to mutualists. Plant J 93:166–180. 10.1111/tpj.13741.

Svozil, J., Gruissem, W., and Baerenfaller, K. (2016). Meselect - A Rapid and Effective Method for the Separation of the Main Leaf Tissue Types. Front Plant Sci 7:1701. 10.3389/fpls.2016.01701.

Tetyuk, O., Benning, U.F., and Hoffmann-Benning, S. (2013). Collection and analysis of Arabidopsis phloem exudates using the EDTA-facilitated Method. J Vis Exp:e51111. 10.3791/51111.

Todesco, M., Rubio-Somoza, I., Paz-Ares, J., and Weigel, D. (2010). A collection of target mimics for comprehensive analysis of microRNA function in Arabidopsis thaliana. PLoS Genet 6:e1001031. 10.1371/journal.pgen.1001031.

Tovar-Aguilar, A., Grimanelli, D., Acosta-Garcia, G., Vielle-Calzada, J.P., Badillo-Corona, J.A., and Duran-Figueroa, N. (2024). The miRNA822 loaded by ARGONAUTE9 modulates the monosporic female gametogenesis in Arabidopsis thaliana. Plant Reprod 37:243–258. 10.1007/s00497-023-00487-2.

Tsikou, D., Yan, Z., Holt, D.B., Abel, N.B., Reid, D.E., Madsen, L.H., Bhasin, H., Sexauer, M., Stougaard, J., and Markmann, K. (2018). Systemic control of legume susceptibility to rhizobial infection by a mobile microRNA. Science 362:233–236. 10.1126/science.aat6907.

Varkonyi-Gasic, E., Wu, R., Wood, M., Walton, E.F., and Hellens, R.P. (2007). Protocol: a highly sensitive RT-PCR method for detection and quantification of microRNAs. Plant Methods 3:12. 10.1186/1746-4811-3-12.

Vlot, A.C., Sales, J.H., Lenk, M., Bauer, K., Brambilla, A., Sommer, A., Chen, Y., Wenig, M., and Nayem, S. (2020). Systemic propagation of immunity in plants. New Phytol 10.1111/nph.16953.

Voinnet, O. (2022). Revisiting small RNA movement in plants. Nat Rev Mol Cell Biol 23:163–164. 10.1038/s41580-022-00455-0.

Voinnet, O. (2025). Three decades of mobile RNA silencing within plants: what have we learnt? J Exp Bot 10.1093/jxb/eraf312.

Weralupitiya, C., Eccersall, S., and Meisrimler, C.N. (2024). Shared signals, different fates: Calcium and ROS in plant PRR and NLR immunity. Cell Rep 43:114910. 10.1016/j.celrep.2024.114910.

Xu, P., Ma, W., Liu, J., Hu, J., and Cai, W. (2021). Overexpression of a small GTP-binding protein Ran1 in Arabidopsis leads to promoted elongation growth and enhanced disease resistance against P. syringae DC3000. Plant J 108:977–991. 10.1111/tpj.15445.

Yildiz, I., Mantz, M., Hartmann, M., Zeier, T., Kessel, J., Thurow, C., Gatz, C., Petzsch, P., Kohrer, K., and Zeier, J. (2021). The mobile SAR signal N-hydroxypipecolic acid induces NPR1-dependent transcriptional reprogramming and immune priming. Plant Physiol 186:1679–1705. 10.1093/plphys/kiab166.

Zhang, Q., Zhao, F., Wu, Z., and Zhu, D. (2022). A simple and robust method for isolating and analyzing chromatin-bound RNAs in Arabidopsis. Plant Methods 18:135. 10.1186/s13007-022-00967-y.

Zhang, W., Gao, S., Zhou, X., Xia, J., Chellappan, P., Zhou, X., Zhang, X., and Jin, H. (2010). Multiple distinct small RNAs originate from the same microRNA precursors. Genome Biol 11:R81. 10.1186/gb-2010-11-8-r81.

Zhang, W., Gao, S., Zhou, X., Chellappan, P., Chen, Z., Zhou, X., Zhang, X., Fromuth, N., Coutino, G., Coffey, M., et al. (2011a). Bacteria-responsive microRNAs regulate plant innate immunity by modulating plant hormone networks. Plant Mol Biol 75:93–105. 10.1007/s11103-010-9710-8.

Zhang, X., Zhao, H., Gao, S., Wang, W.C., Katiyar-Agarwal, S., Huang, H.D., Raikhel, N., and Jin, H. (2011b). Arabidopsis Argonaute 2 regulates innate immunity via miRNA393( *)-mediated silencing of a Golgi-localized SNARE gene, MEMB12. Mol Cell 42:356–366. 10.1016/j.molcel.2011.04.010.

Zhu, J., Lolle, S., Tang, A., Guel, B., Kvitko, B., Cole, B., and Coaker, G. (2023). Single-cell profiling of Arabidopsis leaves to Pseudomonas syringae infection. Cell Rep 42:112676. 10.1016/j.celrep.2023.112676.

