## Supplemental data for "Systemic resistance to pathogens in Arabidopsis requires HASTY-dependent miRNA cell-to-cell movement"

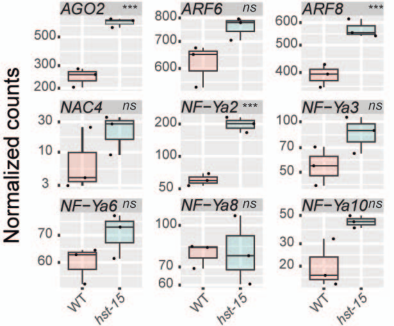


**Figure S1: miRNAs and defense-related genes misregulated in *hst-15* and in samples with systemically induced defenses.**

Box plot showing normalized counts of genes targeted by the six miRNAs labeled in Fig. 1H, in non-treated *hst-15* plants. Significant differences are based on DESeq2 analysis.

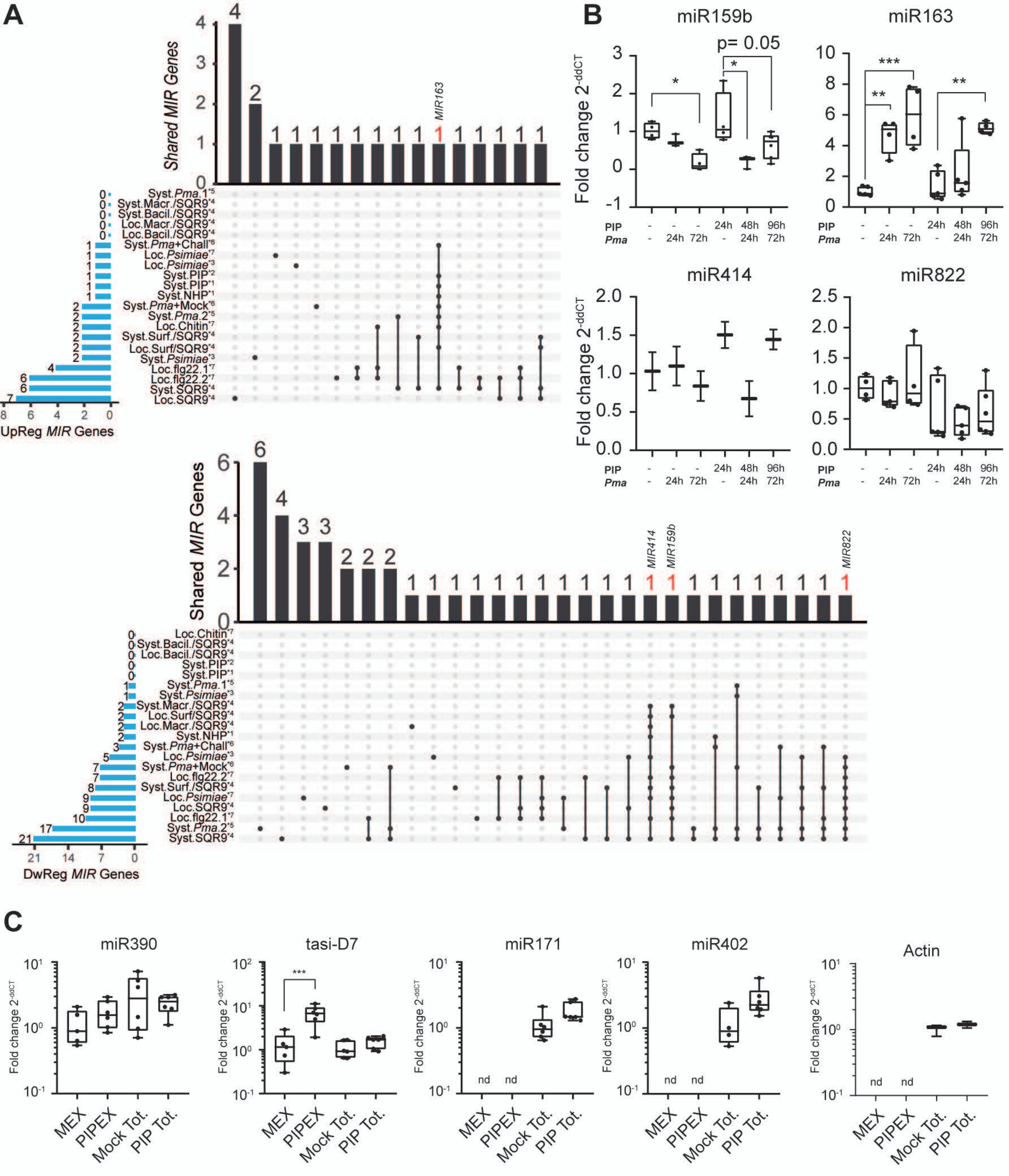


**Figure S2: Selection of miRNAs involved in local and systemic defense activation.**
(A) Upset plot showing pri-miRNAs that are upregulated (top) or downregulated (bottom) in samples treated locally or systemically with different defense inducers. *MIR* genes misregulated in more than nine transcriptomes are highlighted. (B) miRNA abundance in leaves untreated (-), treated systemically with PIP for 24 h, locally infected with *Pma* for 24 or 48 h, or co-treated with PIP and *Pma* (48 h with PIP and by 24 h with *Pma*, or 96 h with PIP and 72 h with *Pma*). (C) miRNA abundance measured by RT-qPCR in phloem exudates collected from Mock-treated (MEX), or PIP-treated (PIPEX) leaves. *: $p < 0.05$, **: $p < 0.01$, ***: $p < 0.001$ by ANOVA with multiple comparisons corrected using Dunn's test (B) or by two-tailed unpaired t-test (C).

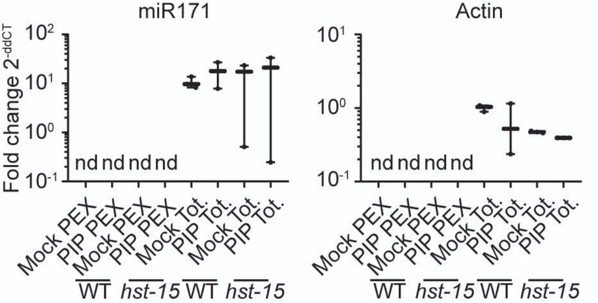


**Figure S3: Detection of miR171 and Actin in phloem exudates.**

Abundance of miR171 and Actin in phloem exudates collected from Mock-treated (MEX) or PIP-treated (PIPEX) leaves of wild-type (WT) and *hst-15* plants. These are negative controls for Figure 5B. *: $p < 0.05$ by ANOVA with multiple comparisons corrected using Dunn's test.

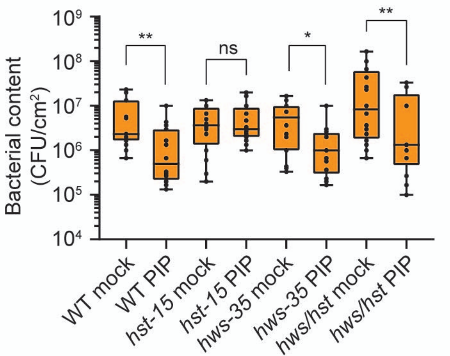


**Figure S4: Pathogen growth in *hst-15* plants complemented with the *hws-35* mutation.** *Pma* growth in leaves of wild-type (WT), *hst-15*, and *hws-35/hst-15* (*hws/hst*) plants plants pre-treated systemically with mock or PIP before pathogen inoculation. Pathogen growth is expressed as CFU/cm². Statistical significance: *: $p < 0.05$, **: $p < 0.01$, determined by Mann–Whitney test.

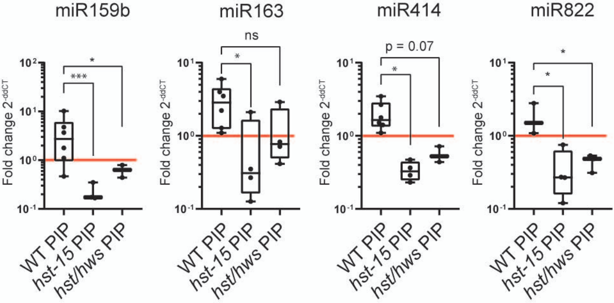


**Figure S5: Restoration of miRNA phloem loading in *hws35/hst-15* double mutant.**

Abundance of miRNAs in phloem exudates collected from Mock-treated (MEX) or PIP-treated (PIPEX) leaves of wild-type (WT), *hst-15*, and *hws-35/hst-15* (*hws/hst*) double mutant plants. Fold change was calculated comparing PIP-treated to Mock-treated samples (indicated by the red line) for each genotype.

*: $p < 0.05$, **: $p < 0.01$, ***: $p < 0.001$ by Kruskal–Wallis test with multiple comparisons corrected using Dunn's test.

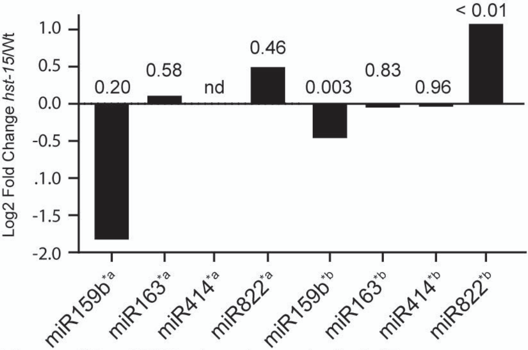


**Figure S6. miRNA abundance in *hst-15*.**
Abundance of miR159b, miR163, miR414, and miR822 in *hst-15* plants from Cambiagno et al 2021 (*a), or Gonzalo et al., 2025 (*b). Fold change and False Discovery Rate (FDR) values are indicated above each bar.

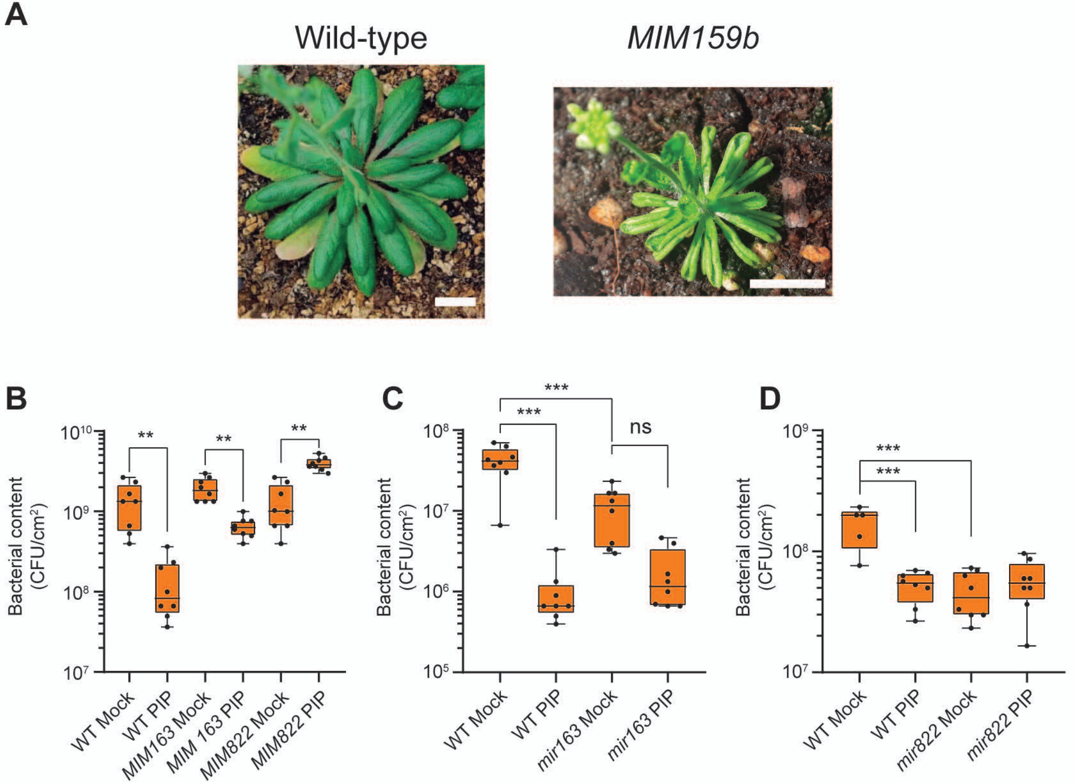


**Figure S7. Pathogen growth in plants with impaired miR163 or miR822 activity.**
(A) Phenotype of 45-day-old wild-type and *MIM159b* plants. White bars represent 0.5 cm. (B) *Pma* growth in leaves of wild-type (WT), *MIM163*, *MIM822*, *mir163*, and *mir822* plants pre-treated systemically with mock or PIP before pathogen inoculation. Bacterial content is expressed as CFU/cm². Statistical significance: *: $p < 0.05$, **: $p < 0.01$, ***: $p < 0.001$, determined by ANOVA with multiple comparisons corrected using Dunn's test.

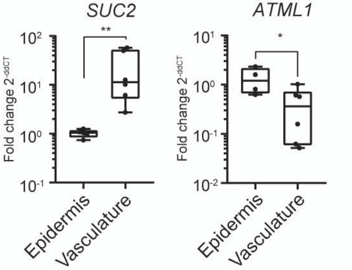


**Figure S8. Enrichment of vasculature and epidermal cell.** Transcript levels of *SUC2* and *ATML1* in vascular- and epidermal-enriched tissues. *: $p < 0.05$, **: $p < 0.01$ by ANOVA test with multiple comparisons corrected using Dunn's test.

**Table S1: Wild-type and mutant plants used in this study.**

| Seeds | Code | Mutation | Transgenes | From |
|---|---|---|---|---|
| **Col-0** | CS22681 | - | - | Manavella´s Lab |
| ***hst-15*** | SALK_079290 | - | - | Manavella´s Lab |
| ***hws-35*** | - | EMS G238R | - | Manavella´s Lab |
| ***hst-15hws-35*** | SALK_079290 | EMS G238R | - | Manavella´s Lab |
| ***hst-3*** | CS24278 | - | - | Manavella´s Lab |
| ***hst-1*** | CS3810 | - | - | Voinnet Lab |
| ***ran1-1*** | SALK_138680 | - | - | Manavella´s Lab |
| ***ran1-2*** | SALK_067649 | - | - | Manavella´s Lab |
| ***impa2*** | SALK_099707 | - | - | Manavella´s Lab |
| ***mir163*** | SALK_034556C | - | - | ABRC |
| ***mir822*** | SALK_023928 | - | - | ABRC |
| **Col-0** | CS22681 | - | *pSUC2:amiRSUL* | Voinnet Lab |
| **Col-0** | CS22681 | - | 35S:MIM163 | In this study |
| **Col-0** | CS22681 | - | 35S:MIM822 | In this study |
| **Col-0** | CS22681 | - | 35S:GFP:HST | Manavella´s Lab |
| **Col-0** | CS22681 | - | pSUC2:HST:GFP | In this study |

**Table S2: Primers used in this study.**

| Oligo for PCR | Strand | Sequence | Use |
|---|---|---|---|
| **Universal Stem-Loop** | RV | GTGCAGGGTCCGAGGT | qPCR |
| **miR159b -Stemloop** | RV | GTCGTATCCAGTGCAGGGTCCGAGGTATTCGCACTGGATACGACaagagc | RT |
| **miR159b** | FW | GCATGCTtttggattgaaggga | qPCR |
| **miR159 stemloop** | FW | AGAAAAGCTGCTAAGCTATGGATCCC | qPCR from Chromatin and Nucleoplasmic enriched samples |
| **miR159 stemloop** | RV | GCCATTAAAGGGCAAGTTAAAGCTC | qPCR from Chromatin and Nucleoplasmic enriched samples |
| **miR159 surrounding the DCL cleavage site** | FW | CGATAGATCTTGATCTGACGATGG | qPCR from Chromatin and Nucleoplasmic enriched samples |
| **miR159 surrounding the DCL cleavage site** | RV | TGGGATCCATAGCTTAGCAGC | qPCR from Chromatin and Nucleoplasmic enriched samples |
| **miR163 -Stemloop** | RV | GTCGTATCCAGTGCAGGGTCCGAGGTATTCGCACTGGATACGACatcgaag | RT |
| **miR163** | FW | GCATGCTttgaagaggacttggaa | qPCR |
| **miR163 stemloop** | FW | GTTCCCGGTTCCTGAGAGTG | qPCR from Chromatin and Nucleoplasmic enriched samples |
| **miR163 stemloop** | RV | TTCTTCGACCGTGCTCTTCC | qPCR from Chromatin and Nucleoplasmic enriched samples |
| **miR163 surrounding the DCL cleavage site** | FW | ACCGACCAAAACCCGGTGG | qPCR from Chromatin and Nucleoplasmic enriched samples |
| **miR163 surrounding the DCL cleavage site** | RV | GAAGTTGTTCCTGGAAGAGAGTGTTG | qPCR from Chromatin and Nucleoplasmic enriched samples |

| | | | |
|---|---|---|---|
| **miR414 - Stemloop** | RV | GTCGTATCCAGTGCAGGGTCCGAGGTATTCGCACTGGATACGACtgacgat | RT |
| **miR414** | FW | GCATGCTtcatcttcatcatc | qPCR |
| **miR822 -Stemloop** | RV | GTCGTATCCAGTGCAGGGTCCGAGGTATTCGCACTGGATACGACcatgtg | RT |
| **miR822** | FW | GCATGCTtgcgggaagcatttg | qPCR |
| **miR822 stemloop** | FW | GGAGAATGAAATCACATTCCATAC | qPCR from Chromatin and Nucleoplasmic enriched samples |
| **miR822 stemloop** | RV | ACCTTCATAGATAGCACATGCTTG | qPCR from Chromatin and Nucleoplasmic enriched samples |
| **miR822 surrounding the DCL cleavage site** | FW | CGCATGTTGTTTTCTGCGGGA | qPCR from Chromatin and Nucleoplasmic enriched samples |
| **miR822 surrounding the DCL cleavage site** | RV | ACCTTCATAGATAGCACATGCTTG | qPCR from Chromatin and Nucleoplasmic enriched samples |
| **miR171c-Stemloop** | RV | GTCGTATCCAGTGCAGGGTCCGAGGTATTCGCACTGGATACGACcgtgatat | RT |
| **miR171c-Stemloop** | FW | GCATGCTTGAGCCGTGCCAAT | qPCR |
| **miR171c stemloop** | FW | CAATCagaaaaccgtactcttttg | qPCR from Chromatin and Nucleoplasmic enriched samples |
| **miR171c stemloop** | RV | CTCAAtcaaataaaccgatcttta | qPCR from Chromatin and Nucleoplasmic enriched samples |
| **miR171c surrounding the DCL cleavage site** | FW | tgagcgcactatcggacatca | qPCR from Chromatin and Nucleoplasmic enriched samples |
| **miR171c surrounding the DCL cleavage site** | RV | caaaagagtacggttttctGATTG | qPCR from Chromatin and Nucleoplasmic enriched samples |
| **miR402-Stemloop** | RV | GTCGTATCCAGTGCAGGGTCCGAGGTATTCGCACTGGATACGACcagaggtt | RT |
| **miR402** | FW | GCATGCTttcgaggcctatt | qPCR |

| | | | |
|---|---|---|---|
| **miR390-Stemloop** | RV | GTCGTATCCAGTGCAGGGTCCGAGGTATTCGCACTGGATACGACggcgct | RT |
| **miR390** | FW | GCATGCTAAGCTCAGGAGGGAT | qPCR |
| **Tas3a-D7-Stemloop** | RV | GTCGTATCCAGTGCAGGGTCCGAGGTATTCGCACTGGATACGACtggggt | RT |
| **Tas3a-D7-Stemloop** | FW | GCGGCGTCTTGACCTTGTAAG | qPCR |
| **RAN1** | FW | GAAGAACAGGCAAGTGAAGGC | qPCR |
| **RAN1** | RV | TCCAGCCAGTTTTCTAGCAAGG | qPCR |
| **IMPA2** | FW | AAGAAGCTGCGTGGGCAATATC | qPCR |
| **IMPA2** | RV | GTGGTACGGCTGCATCATTACC | qPCR |
| **SUC2** | FW | TAGCCATTGTCGTCCCTCA | qPCR |
| **SUC2** | RV | CCTAACACAAATGCTGGAATGT | qPCR |
| **ATML1** | FW | GAGGAGGAGGAGGTAGTGCT | qPCR |
| **ATML1** | RV | TGTGAGTAGTGAACCGCCAC | qPCR |
| **Actin 2/8** | FW | ggTAACATTgTgCTCAgTggTgg | qPCR |
| **Actin 2/8** | RV | GGAGATCCACATCTGCTGGAATG | qPCR |
| **UBC9** | FW | TGGCTTCGAAAAGGATCTTG | qPCR |
| **UBC9** | RV | TCGATATGGTGAGTGCAGGA | qPCR |
| **IPS1** | FW | caccacaaaaacaaaagaaaaatg | cloning mimicry generation |
| **IPS1** | RV | aagaggaattcactataaagag | cloning mimicry generation |
| **IPS1-MIM163** | RV | TCGAAGTTCCAATCAATCCTCTTCAAagcttcggttcccctcg | cloning mimicry generation |
| **IPS1-MIM163** | FW | TGAAGAGGATTGATTGGAACTTCGATtttctagagggagataa | cloning mimicry generation |
| **IPS1-MIM822** | RV | aaCATGTGCAAAactaGCTTCCCGCAagcttcggttcccctcg | cloning mimicry generation |
| **IPS1-MIM822** | FW | ctTGCGGGAAGCtagtTTTGCACATGtttctagagggagataa | cloning mimicry generation |
